# Robust Inference of Population Size Histories from Genomic Sequencing Data

**DOI:** 10.1101/2021.05.22.445274

**Authors:** Gautam Upadhya, Matthias Steinrücken

## Abstract

Unraveling the complex demographic histories of natural populations is a central problem in population genetics. Understanding past demographic events is of general anthropological interest, but is also an important step in establishing accurate null models when identifying adaptive or disease-associated genetic variation. An important class of tools for inferring past population size changes from genomic sequence data are Coalescent Hidden Markov Models (CHMMs). These models make efficient use of the linkage information in population genomic datasets by using the local genealogies relating sampled individuals as latent states that evolve along the chromosome in an HMM framework. Extending these models to large sample sizes is challenging, since the number of possible latent states increases rapidly.

Here, we present our method CHIMP (**C**HMM **H**istory-**I**nference **M**aximum-Likelihood **P**rocedure), a novel CHMM method for inferring the size history of a population. It can be applied to large samples (hundreds of haplotypes) and only requires unphased genomes as input. The two implementations of CHIMP that we present here use either the height of the genealogical tree (*T*_*MRCA*_) or the total branch length, respectively, as the latent variable at each position in the genome. The requisite transition and emission probabilities are obtained by numerically solving certain systems of differential equations derived from the ancestral process with recombination. The parameters of the population size history are subsequently inferred using an Expectation-Maximization algorithm. In addition, we implement a composite likelihood scheme to allow the method to scale to large sample sizes.

We demonstrate the efficiency and accuracy of our method in a variety of benchmark tests using simulated data and present comparisons to other state-of-the-art methods. Specifically, our implementation using *T*_*MRCA*_ as the latent variable shows comparable performance and provides accurate estimates of effective population sizes in intermediate and ancient times. Our method is agnostic to the phasing of the data, which makes it a promising alternative in scenarios where high quality data is not available, and has potential applications for pseudo-haploid data.

**Author Summary:** The demograpic history of natural populations shapes their genetic variation. The genomes of contemporary individuals can thus be used to unravel past migration events and population size changes, which is of anthropological interest. However, it is also important to uncover these past events for studies investigating disease related genetic variation, since past demographic events can confound such analyses. Here we present a novel method for inferring the size history of a given population from full-genome sequencing data of contemporary individuals. Our method is based on a Coalescent Hidden Markov model framework, a model frequently applied to this type of inference. A key component of the model is the representation of unobserved local genealogical relationships among the sampled individuals as latent states. This is achieved by numerically solving certain differential equations that describe the distributions of these quantities and ultimately enables inference of past population size changes. Other methods performing similar inference rely on availability of high quality genomic data, whereas we demonstrate that our method can be applied in situations with limited data quality.

## 1 Introduction

Advances in technology for full-genome sequencing have made it possible to collect large amounts of genomic data for increasingly large samples from many species and diverse population groups. This wealth of genetic data has much to tell us about underlying biological and population genetic phenomena. These datasets can be used to study the relatedness among individuals to study complex demographic histories, helping unravel population size changes, population structure, and migration events. In addition, adaptation of beneficial alleles or other forms of selection leave characteristic signatures in genomic sequencing data. Thus, genetic variation across individuals, in human populations in particular, can be used to reveal genetic factors underlying traits relevant for medical and health-related applications. Genome wide association studies are a widely-used tool to detect such associations. However, effects of population structure can confound the results of these association studies, see for example Barton et al. (2019). Thus, it is imperative to develop population genetic tools for the analysis of whole-genome sequencing data that can infer the underlying demographic history and establish appropriate null models for studying adaptation and associations of genetic variation with certain phenotypes or disease outcomes.

To date, many methods have been presented in the literature that infer different aspects of demographic histories from different signals in the data. The focus in this study is on the inference of the size history of a single population, and here we briefly review methods with a similar focus. Several methods perform inference using the site frequency spectrum (SFS), either assuming no linkage between sites (Liu and Fu, 2015; Bhaskar et al., 2015), or complete linkage (Palacios et al., 2019). These methods can be efficiently applied to large sample sizes which particularly improves their ability to infer recent population size changes (Bhaskar et al., 2015; Keinan and Clark, 2012). However, these methods do not leverage information about decay of linkage disequilibrium along the chromosome, which has been shown to increase power, see for example Figure 2 of Terhorst et al. (2017). Other methods make use of linkage information by fitting demographic models to the empirical distribution of long shared tracts of Identity-By-Descent directly (Browning and Browning, 2015; Palamara et al., 2012). Since these methods consider tracts above a certain length threshold, they are most powerful at inferring recent population size changes. While these methods account for some linkage information, they do not model the correlation in tract length along the genome. Some recent methods aim to directly reconstruct the multi-locus genealogy relating the sampled individuals from high-quality genomic sequencing data (Rasmussen et al., 2014; Kelleher et al., 2019; Speidel et al., 2019). Such genealogies are useful for a variety of down-stream analyses and can be used for demographic inference as well.

A powerful class of methods to infer population size histories that account for linkage, both in terms of length of shared haplotypes and correlation along the genome, are Coalescent Hidden Markov Models (CHMMs). These methods are based on the Sequentially Markovian Coalescent (SMC) introduced by McVean and Cardin (2005), based on work by Wiuf and Hein (1999). In this framework, the correlations among the marginal genealogies relating the sampled individuals at each locus in the genome due to chromosomal linkage and ancestral recombination events is approximated by a Markov chain. The observed genetic variation is subsequently modeled by a mutation process on these marginal genealogies. Using the full marginal genealogies as latent states in a Hidden Markov Model (HMM) framework is prohibitive, but employing lower-domensional summaries of these genealogies facilitates computationally efficient inference of population size histories.

A number of different CHMM-based inference tools have been developed, including `PSMC` (Li and Durbin, 2011), `MSMC` (Schiffels and Durbin, 2014), `MSMC2` (Wang et al., 2020), `SMC++` (Terhorst et al., 2017), and `diCal` (Sheehan et al., 2013; Steinrücken et al., 2019). These methods differ in the sample size that they can analyze and in how the marginal genealogies are represented in the respective CHMM. For example, `PSMC` can only be applied to samples of size 2, whereas `MSMC2` is commonly applied to samples with sizes around 10. However, the computational cost of the latter does increase substantially with sample size. `SMC++` can be applied to large samples and the data does not need to be phased, whereas `diCal` requires phased data and is also only applicable to sample sizes around 10. The specific implementation details result in each method performing well for certain sample sizes and for certain time periods (Spence et al., 2018; Sellinger et al., 2021), but no method performs uniformly well across all parameter regimes.

Here, we present our novel CHMM method, `CHIMP` (**C**HMM **H**istory-**I**nference **M**L **P**rocedure). We present two implementations of `CHIMP` that differ in the hidden state space that they use for the CHMM. One implementation uses the *T*_*MRCA*_, the time to most recent common ancestor of the local genealogical tree, while the other uses, *ℒ*the total branch length of the tree. Our method uses the number of derived alleles at a given site as the emission of the HMM, and is therefore agnostic to the phasing of data. Moreover, we implemented a flexible composite likelihood scheme that enables efficient scaling to large sample sizes, resulting in runtimes faster than `MSMC2`, specifically for the implementation using *T*_*MRCA*_. The latter also shows comparable inference accuracy in intermediate times, around the Out-Of-Africa bottleneck and more recently in humans, and outperforms other methods for ancient times. Since the method is agnostic to phasing, it has potential applications to pseudo-haploid data.

The paper is organized as follows. In Section 2, we present the general SMC framework that is the basis for CHMMs, and detail the implementation steps for our specific choice of the latent variables. Extending previous work by Miroshnikov and Steinrücken (2017), we present an algorithm to efficiently compute the necessary transition and emission probabilities for the CHMM by numerically solving certain systems of differential equations and incorporate them into a standard Expectation-Maximization (EM) framework for maximum-likelihood inference. In Section 3, we compare the performance of `CHIMP` to other state-of-the-art methods, specifically `MSMC2` (Wang et al., 2020) and `Relate` (Speidel et al., 2019), in several simulation studies over a range of demographic scenarios, and also present an application of our method to data from the 1000 Genomes dataset. Lastly, in Section 4, we discuss possible extensions of our framework to infer more complex demographic histories involving multiple populations and to analyze time stratified samples characteristic of ancient DNA datasets. We also discuss how the posterior distribution of the latent states could be applied to study signatures of selection in the genome.

## 2 Novel CHMM Methods

In this section, we will present relevant background on the Sequentially Markovian Coalescent (SMC) which is the basis for our method, and we will describe our implementation of an HMM framework for inference of past population sizes using *T*_MRCA_ or the total branch length *ℒ* as the hidden states.

### 2.1 CHMM Model under Variable Population Size

The genetic variation observed in a sample of *n* haploid sequences from a given population is affected by its population size history *N* (*k*), where *N* (*k*) is the number of diploid individuals in the population *k* generations before present. We use coalescent theory to model the effects that a time varying population size has on the genealogy of the sample, which in turn affects the pattern of observed genetic variation. In the coalescent framework, it is convenient to measure time in units of 2*N* (0) generations and to consider the population size relative to the size at present. To this end, we introduce the relative coalescent-scaled population size

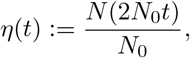

where *N*_0_ ≡ *N* (0) is an arbitrarily chosen reference population size and *k* = 2*N*_0_*t*.

The single-locus coalescent models the genetic variation among *n* sampled haploids at a particular locus in the genome (Kingman, 1982). In the coalescent, the genealogy of the sample is described by following the ancestral lineages of the *n* haplotypes (sampled at present) back in time. Each pair of lineages can coalesce (find a common ancestor) at a given rate *λ*(*t*) that can vary with time *t*. The coalescent rate is the inverse of the relative population size *λ*(*t*) = 1*/η*(*t*), which reflects the fact that ancestral lineages coalesce faster in small populations but coalescence is slower when the population size is large. This process proceeds until all lineages coalesce into a single lineage, referred to as the most recent common ancestor (MRCA), the genetic ancestor of all haplotypes in the sample. The time of this final coalescent event is denoted by *T*_MRCA_. The coalescent thus gives the distribution of genealogies at a single locus. One can model the observed genetic variation at the given locus by superimposing mutations on the genealogy according to a Poisson process with rate *θ/*2, where *θ* = 4*N*_0_*μ* is the population-scaled mutation parameter and *μ* is the per generation per site mutation probability.

The standard coalescent models the marginal genealogy at a single locus. To analyze genomic sequence data, one can use the ancestral recombination graph (ARG), which extends the regular coalescent model to describe the full multi-locus genealogy for *n* sampled haplotypes across *L* loci (Griffiths and Marjoram, 1997; Hudson, 2002). Specifically, the ARG models the genealogies at each individual locus and their correlations induced by presence or absence of ancestral recombination events. Just as in the single-locus case, mutations can be superimposed onto these genealogies to model the observed genetic variation in multi-locus genomic sequence data. While the ARG is a useful tool to simulate genomic data (Hudson, 2002; Kelleher et al., 2016), in many scenarios its applicability in likelihood-based population genetic inference is hindered by its complexity: The space of possible ARGs grows quickly with the number of samples and the length of the genome.

One factor contributing to the complexity of the ARG is the fact that the marginal genealogies at distant loci can depend on each other (Wiuf and Hein, 1999). The Sequentially Markovian Coalescent (SMC), introduced by McVean and Cardin (2005), and its modified version SMC’, introduced by Marjoram and Wall (2006), simplifies the model by assuming that the distribution of the marginal genealogy at a given locus only depends on the genealogy at the previous locus in the sequence, that is, it assumes that the sequence of marginal genealogies is a Markov chain. Under the SMC, the sequence of marginal genealogies is generated as follows. The genealogy at the first locus is distributed according to the standard coalescent. To proceed from one locus to the next, ancestral recombination events occur according to a Poisson process at rate 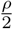 on the branches of the current genealogy, where *ρ* = 4*N* (0)*r*, and *r* is the per base-pair per generation recombination probability. If no recombination events occur, the marginal genealogy is copied unchanged to the next locus. However, if recombination does occur, the lineage above the recombination event is removed up to the next coalescent event involving this lineage. To obtain the genealogy at the next locus, the removed lineage is then replaced by a new lineage that undergoes the standard coalescent dynamic, ie. it can coalesce with the regular coalescent rate into the other ancestral lineages. The distribution of the genealogical trees at each locus is fully determined by the genealogy at the previous locus in the sequence, and thus the sequence of genealogical trees is a Markov chain. An illustration of this generative process for the marginal genealogies is depicted in panels A) and B) of Figure 1.

**Figure 1:**
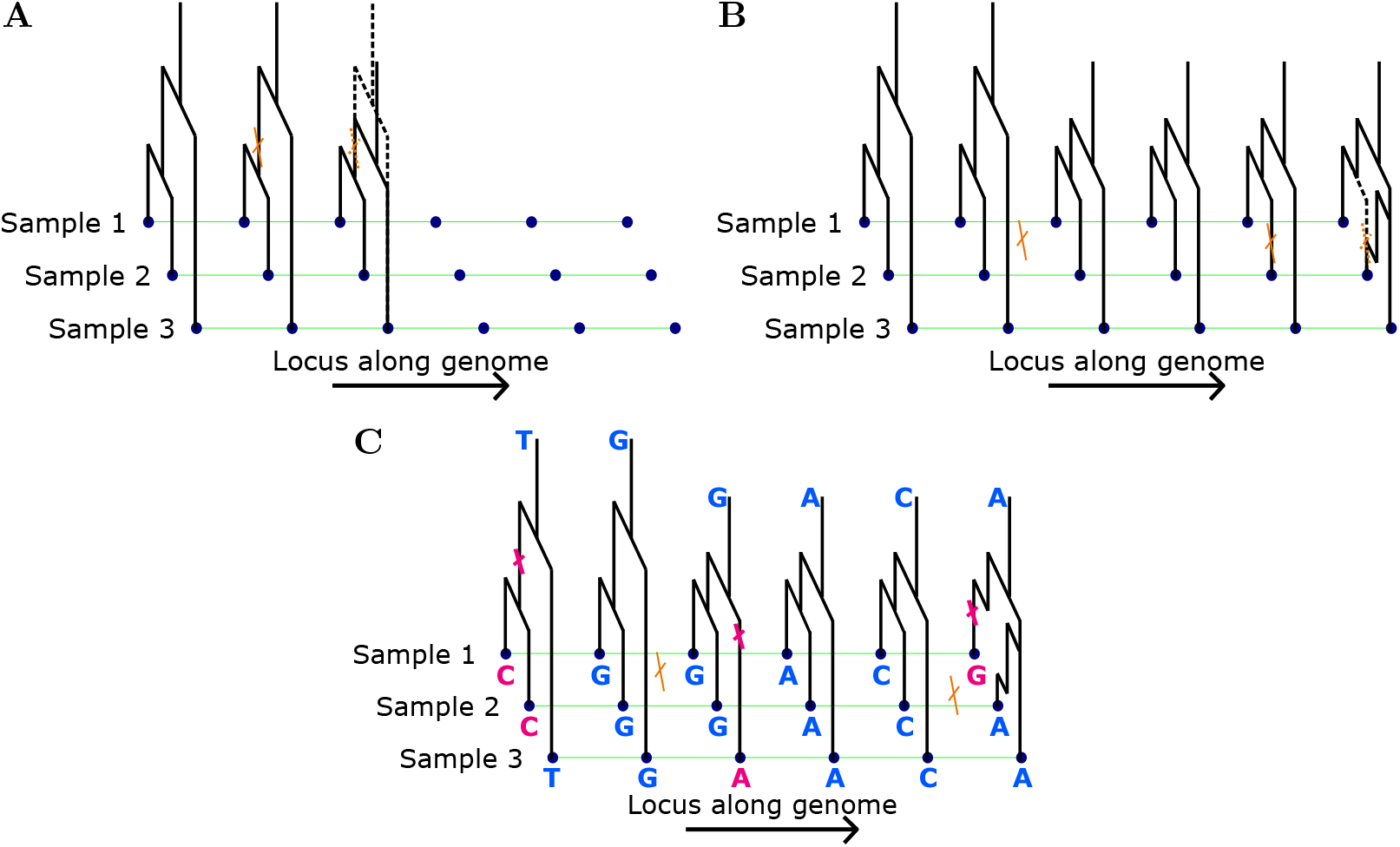
Panel **A** shows the marginal genealogy being propagated unchanged along the genome until an ancestral recombination event is encountered, and the genealogy modified accordingly. In panel **B**, the new genealogy is propagated until a second recombination event is encountered. Panel **C** demonstrates a realization of the mutation process along the genealogy at each locus and the resulting observed genetic data.

**Figure 2:**
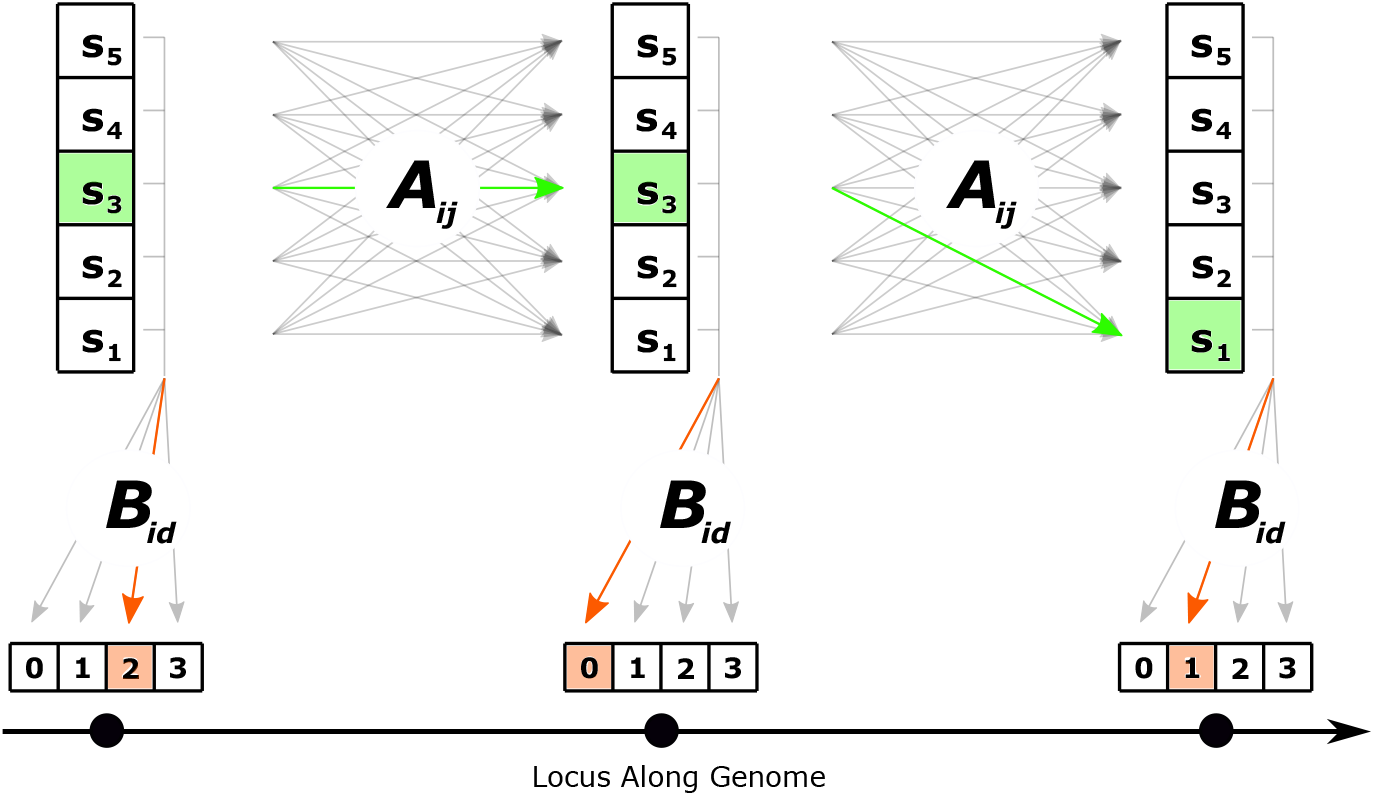
Schematic of our CHMM for a sample of size 4. Information about the underlying tree at each locus is captured by the state *s*_*i*_. 𝒮 = {*s*_1_, …, *s*_5_} is the set of intervals into which the respective summary of the tree (*T*_*MRCA*_ or *ℒ*) can fall. The states change from each locus to the next in accordance with the transition matrix **A** and the observed number of derived alleles at each locus is emitted in accordance with the emission probabilities **B**.

CHMMs use the SMC as the basis for computing likelihoods of observed genomic sequence data. The marginal genealogies are the hidden states in the HMM, and by superimposing mutations onto these trees, the likelihood of the observed genetic variation can be computed as emissions conditional on the hidden state at a given locus (Panel C of Figure 1). However, implementing the full model depicted in Figure 1 in a likelihood-based inference method is intractable due to the prohibitively large hidden state space, which is a consequence of the continuous nature of the genealogical times and the fact that the number of topologies grows super-exponentially with the number of samples. Thus, most existing implementations of CHMMs use a suitable discretization of time and approximate the full local genealogical trees using lower dimensional summaries (often only one-dimensional) to arrive at a finite hidden state space for the HMM. We note that discretizing time is not always necessary when using this framework, as demonstrated by Ki and Terhorst (2020).

We introduce a novel method, `CHIMP`, which is a CHMM with a one-dimensional hidden state space. In our method, we either use the *T*_MRCA_ (time to most recent common ancestor, i.e. tree height) or *L* (total branch length of the tree) as the hidden state. We use *S* +1 increasing times (or lengths) *t*_0_ = 0 *< t*_1_ *<* … *< t*_*S*_ = ∞ to partition the positive real numbers into *S* discrete intervals. The CHMM is in state *s*_*i*_ at locus *ℓ* if *t*_*i−*1_ ≤ *T*_*ℓ*_ *< t*_*i*_, where *T*_*ℓ*_ denotes the *T*_MRCA_ at locus *ℓ* (and likewise for *ℒ* _*ℓ*_). The set of possible states {*s*_1_, …, *s*_*S*_} is denoted by *𝒮*. The sequence of states the CHMM occupies along the complete genome is 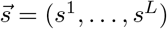, where *s* is the state at locus *ℓ*.

For the emission observed at a given locus, our method uses the number of derived alleles *d* at that locus. Since the data consists of *n* haplotype sequences, we can observe up to *n −* 1 derived alleles at a locus, thus the set of possible emissions is *𝒟* := {0, …, *n −* 1}. Note that *𝒟* includes 0 to model loci where all samples share the same allele. The vector of observations across the genome is 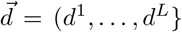, where *d*^*ℓ*^ is the number of derived alleles at locus *ℓ*. With these definitions for the state space and emission space for our CHMM, we introduce the transition and emission probabilities, given by matrices **A** and **B**, respectively, with elements

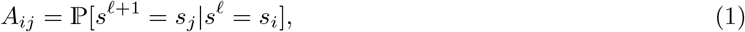

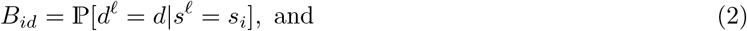

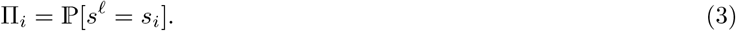

The quantity **Π** is the marginal distribution of the hidden states, and thus it is also the distribution of *s*^0^, the first state in the CHMM. Figure 2 depicts a schematic of the transition and emissions in this CHMM.

We note that if the ancestral or derived state of the alleles is not known, it is possible to instead use the number of minor alleles and adjust the emission probabilities appropriately by folding them. However, our current implementation does not support this. In addition, every locus where all individuals share the same allele is counted as *d* = 0. If the ancestral allele is known, one could in principle distinguish between all individuals sharing the ancestral or the derived allele. In the latter case, the respective mutation would have happened before the MRCA of the sample, which could possibly indicate a more recent MRCA at the respective locus. This scenario could in principle be included in model. However, we do not incorporate this into our model and leave it for future exploration, since it would require assumptions about the divergence times from the outgroups, and it is unclear whether it would improve accuracy of the inference substantially.

### 2.2 T_MRCA_ **as Hidden State**

To use *T*_*MRCA*_ as the hidden state in our CHMM, we discretize the continuous random variable into discrete intervals partitioned at certain *t*_*i*_ as described in Section 2.1. We now describe the numerical methods used to compute the corresponding transition and emission probabilities in equation (1), (2), and (3).

#### 2.2.1 Transition Probabilities

To compute the transition probabilities, we employ an augmented ancestral process with recombination, *𝒜*^*ρ*^, introduced by Miroshnikov and Steinrücken (2017). This process closely resembles the regular ancestral process with recombination described by Simonsen and Churchill (1997) and describes the joint distribution of the genealogies of *n* samples for two adjacent loci separated by a recombination distance of *ρ*. We use *𝒜*^*ρ*^ to compute the respective transition probabilities in the matrix **A**.

The process *𝒜*^*ρ*^(*t*) is initialized at the present (*t* = 0) and tracks the ancestral lineages at two loci, *a* and *b*, simultaneously as they evolve backwards in time. Initially, there are *n* lineages, each ancestral to both loci *a* and *b* of one of the *n* sampled haplotypes. As in the standard coalescent with varying population size, ancestral lineages coalesce with rate *λ*(*t*). Additionally, recombination events can occur on each lineage ancestral to two loci at rate 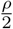 and decouples the dynamics of the two loci. The two decoupled lineages then evolve independently and yield distinct genealogies for each of the loci. Ultimately all lineages coalesce into a common ancestor for both loci.

The states of *𝒜*^*ρ*^(*t*) are denoted by tuples describing the configuration of lineages, (*k*_*ab*_, *k*_*a*_, *k*_*b*_, *κ*). Here, *k*_*ab*_ is the number of active lineages ancestral to both loci (coupled), *k*_*a*_ are the lineages ancestral only to locus *a*, and *k*_*b*_ are the lineages ancestral only to *b* (uncoupled). Finally, *κ* explicitly tracks the number of recombination events that have occurred since *t* = 0. While *k*_*ab*_ ∈ {1, 2, …, *n*}, the decoupled lineages are constrained such that *k*_*a*_, *k*_*b*_ ∈ {0, 1, …, *κ*} since there can be at most as many uncoupled lineages as recombination events. In the full ancestral process, *κ* takes values from 0 to ∞, since there can be an arbitrary number of recombination events between the two loci. The unboundedness of *κ* renders this full process difficult to solve. In the remainder of this work, we restrict *κ* to be at most 1 (and consequently also restrict *k*_*a*_ and *k*_*b*_ to be 0 or 1). This restriction is motivated by the assumption that the recombination rate between two adjacent loci is low, so we expect to see at most one ancestral recombination event separating the genealogies at two neighboring loci. Henceforth, *𝒜*^*ρ*^(*t*) will refer to this restricted process. We speculate on the consequences if this assumption is violated in Section 4.

Figure 3 shows an example trajectory of this augmented ancestral process. Recombination events decouple a shared lineage, while coalescence events fuse two active lineages. If an uncoupled lineage (one of type *k*_*a*_ or type *k*_*b*_) coalesces with a coupled lineage (one of type *k*_*ab*_), the resulting lineage contains ancestry that traces to both loci *a* and *b* in our sample, and is therefore a coupled lineage. We note that *𝒜*^*ρ*^ allows for a joint lineage to recombine (split) and immediately coalesce back together.

**Figure 3:**
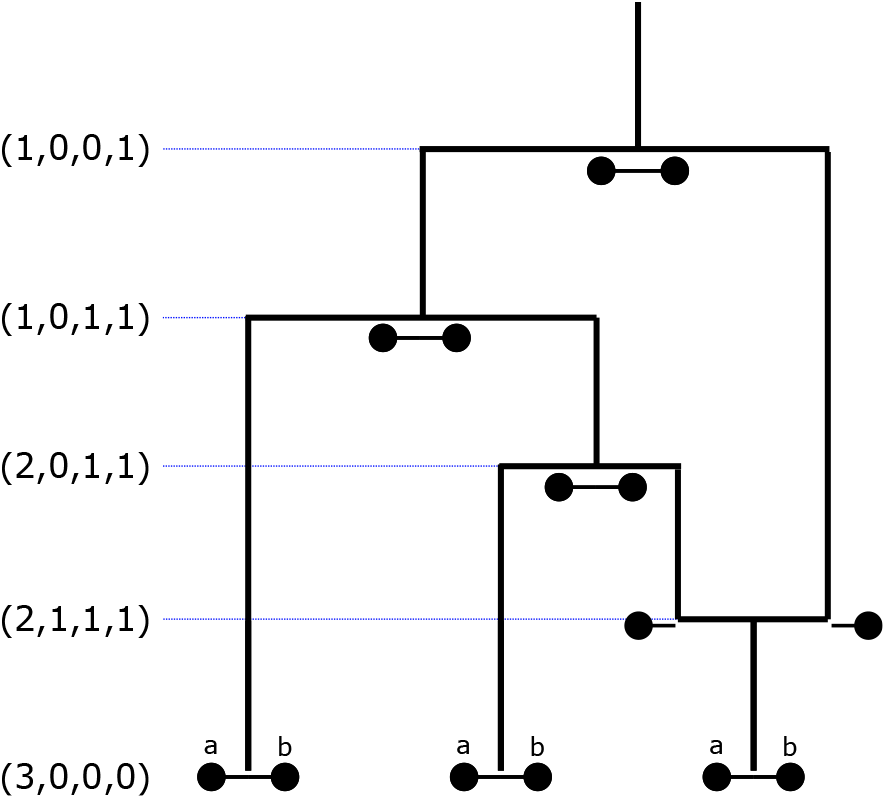
Example trajectory of *𝒜*^*ρ*^(*t*) with the state denoted by tuples. At *t* = 0 there are three lineages of type *k*_*ab*_ ancestral to both loci for their respective samples. The lineages split at ancestral recombination events and join at coalescence events where they find a common ancestor. The trajectory ultimately culminates in the state (1, 0, 0, 1), signifying that there is one lineage ancestral to both loci in all present-day samples, and that one recombination event occurred in this genealogy.

We set *𝒜*^*ρ*^(0) = (*n*, 0, 0, 0), which corresponds to initializing the process in a state with *n* lineages at the present time, each ancestral to both loci in a single sample. The absorbing states are (1, 0, 0, 1) and (1, 0, 0, 0), which correspond to the states where all lineages for both loci have fully coalesced after 0 or 1 recombination events have occurred.

The possible transitions between states and their respective rates are given in Table 1. The rates in the first three rows of this table correspond to all possible coalescent events. These rates are all proportional to the time-dependent coalescent rate *λ*(*t*). The first row describes coalescence among the lineages ancestral to locus *a* and *b*, which results in reducing the number of these lineages by 1. The rate for these events is proportional to all possible pairs of such lineages. The second row describes events that reduce the number of lineages ancestral to only *a* by one, which can either be a coalescent event among the *a* lineages, or a coalescence event between one lineage ancestral to *a* and another ancestral to *a* and *b*. Again, the rate is proportional to the number of such lineage pairings. The third row describes the respective events for the *b* lineages. The fourth row corresponds to recombination events, with a rate proportional to the recombination rate *ρ/*2. These recombination events can only happen in lineages ancestral to *a* and *b* and reduces their number by one. Such events result in one lineage ancestral to only *a* and one only to *b*, increasing the respective numbers by one, and also increasing *κ* by one. Note that the rates for the recombination events reflect the fact we restrict the process to have at most one recombination event.

**Table 1:**
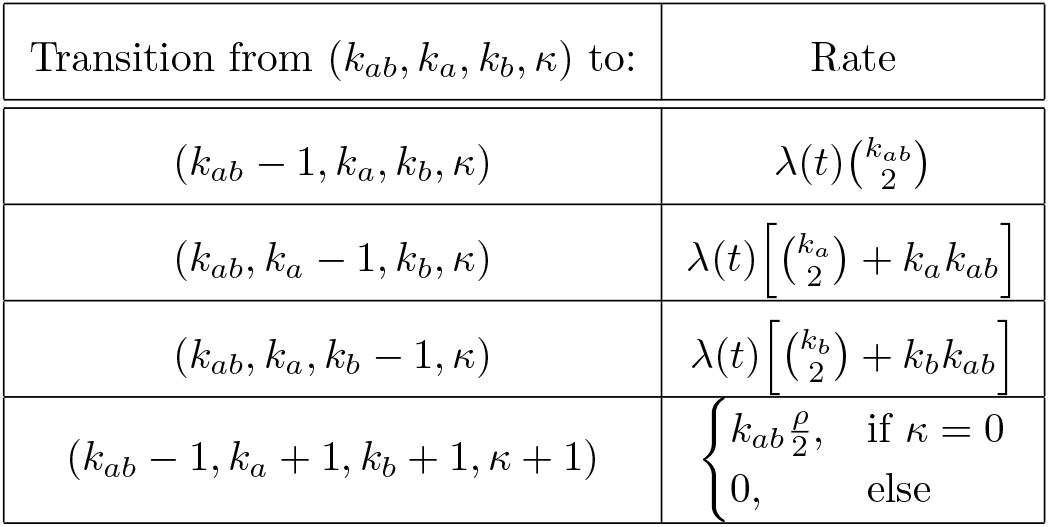
The table shows the possible transitions out of a given state (*k*_*ab*_, *k*_*a*_, *k*_*b*_, *κ*) and their respective rates. The first row gives the rate for coalescence between two lineages that are ancestral to both loci. The second row gives rate for two types of events, coalescences between two lineages ancestral to only locus *a*, and coalescences of a lineage ancestral only to *a* with a lineage ancestral to both. The third row reflects similar events for locus b. The last row gives the rate of recombination events. Note that these rates are defined to permit a maximum of 1 ancestral recombination event occurring between locus *a* and *b*.

| Transition from $(k_{ab}, k_a, k_b, \kappa)$ to: | Rate |
| --- | --- |
| $(k_{ab} - 1, k_a, k_b, \kappa)$ | $\lambda(t) \binom{k_{ab}}{2}$ |
| $(k_{ab}, k_a - 1, k_b, \kappa)$ | $\lambda(t) \left[ \binom{k_a}{2} + k_a k_{ab} \right]$ |
| $(k_{ab}, k_a, k_b - 1, \kappa)$ | $\lambda(t) \left[ \binom{k_b}{2} + k_b k_{ab} \right]$ |
| $(k_{ab} - 1, k_a + 1, k_b + 1, \kappa + 1)$ | $\begin{cases} k_{ab} \frac{\rho}{2}, & \text{if } \kappa = 0 \\ 0, & \text{else} \end{cases}$ |

Define 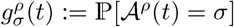 to be the probability that the augmented ancestral process is in state *σ ∈ ℛ* at time *t*, where *ℛ* is the set of all possible states. Then 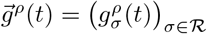 is the distribution of the process at time *t*, a vector of probabilities over all the states *σ ∈ ℛ*. Since *𝒜*^*ρ*^ is a continuous-time Markov process, the evolution of 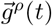 is given by the system of ordinary differential equations (ODEs)

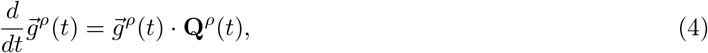

where **Q**^*ρ*^(*t*) is the rate matrix consisting of the rates given in Table 1. The rate matrix is time-dependent, since the coalescent rates *λ*(*t*) are as well. We can now obtain the probabilities 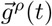 (*t*) by numerically integrating equation (4).

Moreover, from the distribution 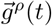, we can compute the cumulative joint distribution function (CDF) of the *T*_MRCA_ at the two loci, ℙ [*T*_*a*_ ≤ *τ*_*a*_; *T*_*b*_ ≤ *τ*_*b*_], where 0 ≤ *τ*_*a*_, *τ*_*b*_ *< ∞*, and *T*_*a*_ and *T*_*b*_ are the *T*_MRCA_’s at *a* and *b* respectively. Without loss of generality, we assume that *τ*_*a*_ *< τ*_*b*_ and obtain

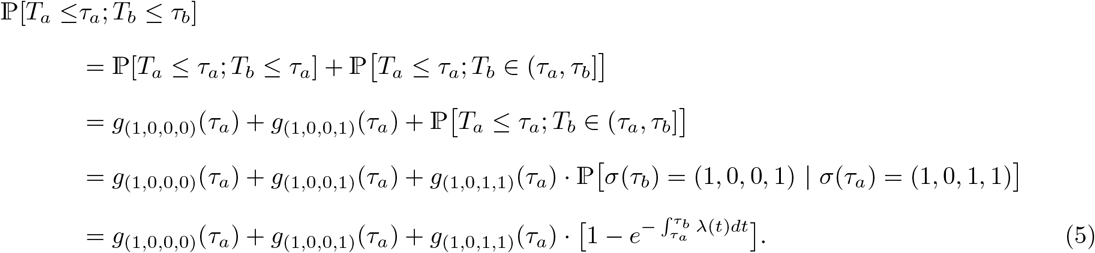

In the first equality, we partition the probability according to whether *T*_*b*_ *< τ*_*a*_ or *T*_*b*_ *> τ*_*a*_. The second equality holds, because ℙ [*T*_*a*_ ≤ *τ*_*a*_; *T*_*b*_ ≤ *τ*_*a*_] is the probability that the ancestral process is in an absorbing state where both loci have found the *T*_*MRCA*_ by time *τ*_*a*_. The third equality follows from the fact that ℙ[*T*_*a*_ ≤ *τ*_*a*_; *T*_*b*_ *∈* (*τ*_*a*_, *τ*_*b*_]] is the probability that the lineages at *a* found a common ancestor by time *τ*_*a*_ and the lineages at *b* find a common ancestor after *τ*_*a*_, but before *τ*_*b*_, which is only possible if a recombination event occurred. Since this term is conditional on *a* having found its *T*_*MRCA*_, only 2 lineages can be remaining (one ancestral only to *b*, and one the common ancestor of *a*) due to the assumption that *κ* ≤ 1, and thus the term simplifies to the coalescence probability of two lineages between times *τ*_*a*_ and *τ*_*b*_. The final equality follows.

By evaluating equation (5) at the values 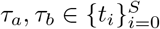, the interval boundaries for the discretized state space, we can obtain a joint cumulative distribution function (CDF) for *T*_*a*_ and *T*_*b*_, denoted **A**^*CDF*^. From this it is straightforward to compute the matrix that comprises the values of the discrete joint probability density function (PDF), **A**^*P DF*^. Dividing the joint probabilities by the marginal probabilities, we arrive at **A**, the transition probability matrix itself:

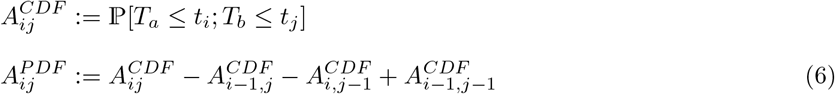

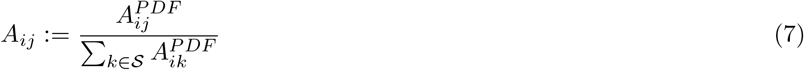

Additionally, we can obtain the vector of marginal probabilities **Π** as

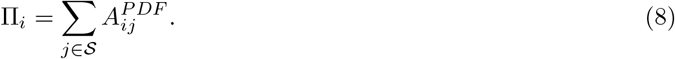

#### 2.2.2 Emission Probabilities

To compute the emission probabilities with *T*_MRCA_ as the hidden state, we introduce an augmented single-locus ancestral process with mutation *𝒜*^*θ*^ which is an extension to the regular ancestral process (Griffiths and Marjoram, 1997) that is motivated by the fact that, conditional on the coalescent tree, mutations are Poisson distributed along the branches with rate 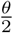. Similar to the recombination case, we only consider at most one mutation event, motivated by the assumption that the per locus mutation rate is low. We speculate on the consequences if this assumption is violated in Section 4. The states of this process are denoted by (*k, k*^∗^), where *k* is the number of active lineages ancestral to the *n* samples, and *k*^∗^ is the number of lineages that were active at the time of the first mutation event along the genealogy (going backwards in time). If no mutation has occurred yet, *k*^∗^ assumes a value of − 1.

The process is initialized in (*n*, − 1), a state before any mutation has occurred with one ancestral lineage for each sample. The transition rates are given in Table 2 and an example trajectory is shown in Figure 4. The possible transitions are either two lineages coalescing or a lineage mutating. The rate for coalescence is given by the coalescent rate *λ*(*t*) times the number of possible pairs that can coalesce, and such an event reduces the number of lineages by one. The rate for a transition via mutation is given by the mutation rate 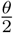 multiplied with the number of lineages that a mutation can occur on. The number of lineages that were active at the time of the mutation event is recorded in the second component of the state. Since we restrict to one mutation event at most, if this number is set once, it will not be set again. The process is absorbed in any state (1, *k*^∗^) with *k*^∗^ *∈* {*−*1, 2, …, *n*}. Note that it is important to continue the process after a mutation event occurred until all lineages are coalesced so that we obtain the full distribution of the *T*_MRCA_.

**Table 2:** The transition rates of the augmented ancestral process *𝒜*^*θ*^. The first row gives the rate of a coalescence event of two lineages, while the second row gives the rate for mutation events. Note that only one mutation event is permitted.

| Transition | Rate |
| --- | --- |
| $(k, k^*) \rightarrow (k-1, k^*)$ | $\lambda(t) \binom{k}{2}$ |
| $(k, -1) \rightarrow (k, k)$ | $k \frac{\theta}{2}$ |

**Figure 4:**
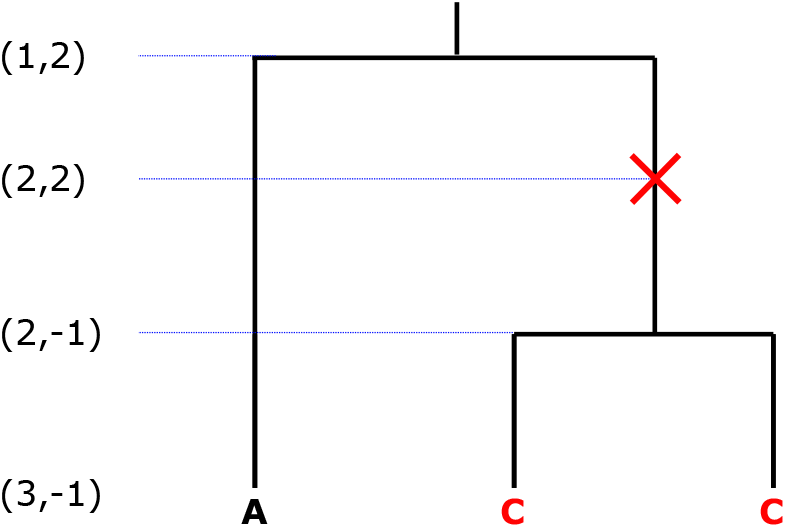
Example trajectory of the ancestral process with mutation for *n* = 3 samples with the state (*k, k*^∗^) indicated on the left. The mutation process is superimposed onto the regular genealogical process. In this example, the mutation happens when there are two ancestral lineages, resulting in two samples carrying the derived allele.

Similar to the procedure for *𝒜*^*ρ*^, we collect all transition rates in a matrix **Q**^*θ*^(*t*). We further define 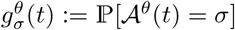 as the probability that the ancestral process *𝒜*^*θ*^ is in a state *σ ∈ ℳ* at time *t*, where *ℳ* is the set of all possible states. The evolution of the vector of all probabilities 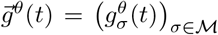 is given by the system of ODEs

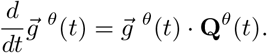

Again, we obtain the solution to these ODEs numerically.

Using the distribution of this process, we can compute the cumulative distribution of *T*_*MRCA*_ jointly with the probability of emitting *d* derived alleles as

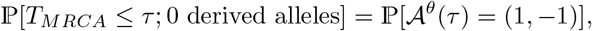

since this gives the probability that all lineages are coalesced by *τ* and no mutation occurred, and

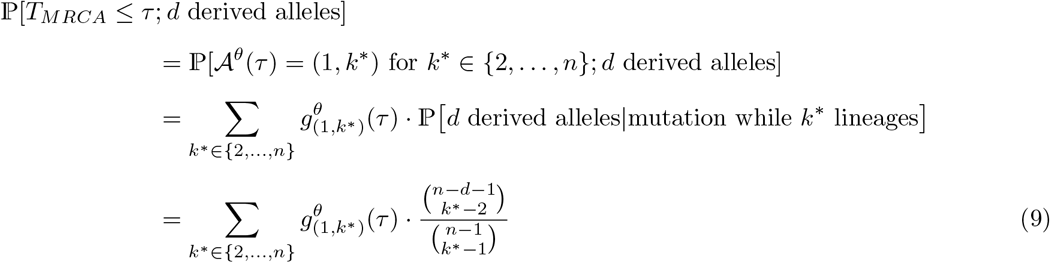

for *d* ϵ{1, …, *n* −1}. The first equality in equation (9) follows from the fact that *T*_*MRCA*_ *< τ* if and only if the ancestral process has found an absorbing state (all lineages have coalesced) before *τ*. In the second equality, we partition this probability with respect to the specific number of lineages active when the mutation occurred, which is encoded in the absorbing state. For the last equality, we substitute the probability of emitting a certain number of derived alleles given that there were *k*^∗^ active lineages at the time of the mutation. This probability is given by the probability that one of the *k*^∗^ lineages subtends *d* leafs (e.g. Durrett, 2008, Ch. 2.1). It is independent of the time of the mutation.

By evaluating these probabilities at times 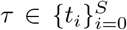, we compute the discretized joint CDF for the emissions, **B**^*CDF*^, which is again used to compute the joint probabilities **B**^*PDF*^ and ultimately the emission probabilities **B** for the CHMM by conditioning on the hidden state:

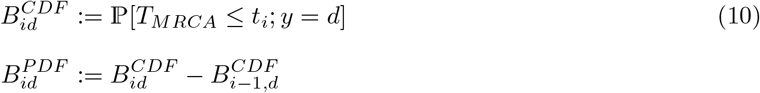

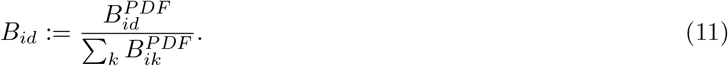

The transition and emission probabilities can then be used to compute likelihoods of observed sequence data and perform inference using an EM algorithm, which will be described in more detail in Section 2.4.

### 2.3 Total Branch Length as Hidden State

We now describe the implementation of our CHMM with the total branch length (sum of all branch lengths) of the genealogical tree *ℒ* as the hidden state at each locus. As before, we discretize *ℒ* by partitioning the real line with a set of values *t*_0_ = 0, *< t*_1_ *<* … *< t*_*S*_ = *∞*.

#### 2.3.1 Transition Probabilities

We use an approach introduced by Miroshnikov and Steinrücken (2017) to compute the joint distribution of the marginal total tree length at locus *a* and *b*. We begin by computing the joint distribution of the total tree length accumulated up to a certain time *t* in the past. Using the augmented ancestral process *𝒜*^*ρ*^ (introduced in Section 2.2.1), which computes the requisite distributions for *T*_*MRCA*_, together with the quantity

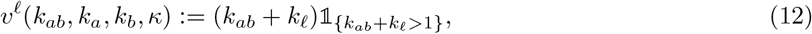

where *ℓ ∈* {*a, b*}, we define and

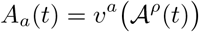

to count the number of active lineages that are ancestral to locus *a* or *b* at a given time *t*. Note that this includes the lineages ancestral to both loci. We define the marginally accumulated tree length

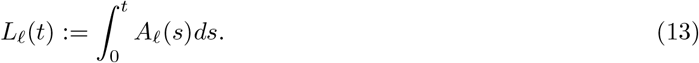

The quantities *L*_*a*_(*t*) and *L*_*b*_(*t*) can be thought of as the total branch lengths that has aggregated at each locus as the process evolves back in time. This holds because the integrand in equation (13) is the number of lineages at a specific locus at a given time and total branch length is accumulated linearly along each active lineage. The indicator function in equation (12) signifies that the process stops accumulating tree length once only a single lineage is left, that is, the *T*_MRCA_ is reached. Using this notation, we now define the probabilities

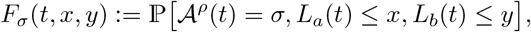

similar to Miroshnikov and Steinrücken (2017), which give the joint distribution of tree length accumulated at both loci up to time *t* and of the ancestral process *𝒜*^*ρ*^(*t*) being in state *σ*.

Miroshnikov and Steinrücken (2017) show that the values of 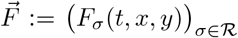 can be obtained as the solutions of the system of partial differential equations (PDEs)

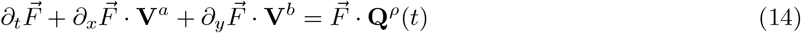

and its corresponding boundary conditions

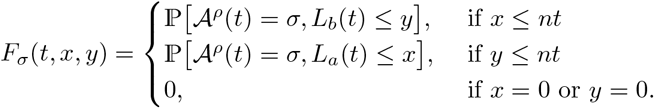

These equations are given in terms of the rate matrix **Q**^*ρ*^(*t*) of the augmented ancestral process *𝒜*^*ρ*^ and the diagonal matrices

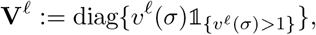

which represent the accumulation of tree length along the active ancestral lineages.

As shown by Miroshnikov and Steinrücken (2017), the quantities ℙ[*𝒜*^*ρ*^(*t*) = *σ, L*_*a*_(*t*) *x* =: *F*_*σ*_(*t, x*) (and the corresponding quantities for *b*) can in turn be obtained as the solution of the PDEs

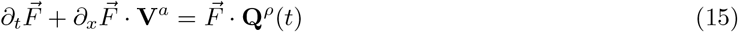

with boundary conditions

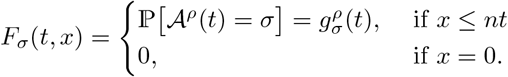

We implemented the scheme introduced in (Miroshnikov and Steinrücken, 2017, Appendix B) to compute the solutions to these PDEs and provide the details of our implementation in Section 1 in S1 Text.

Lastly, the joint distributions of tree length at loci *a* and *b* can be obtained from the solutions of the absorbing states of *𝒜*^*ρ*^ and is given by

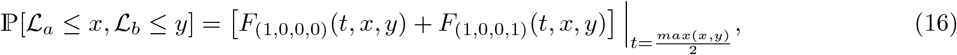

where *ℒ*_*a*_ and *ℒ*_*b*_ denote the total branch length of the genealogies at locus *a* and *b* respectively. Evaluating these probabilities at *x, y* ∈ 𝒮= *t*_0_, *t*_1_, …, *t*_*S*_ yields the elements of the joint cumulative probability matrix **A**^*CDF*^. This discretized joint distribution can then be used in equations (6) and (7) to compute **A**^*PDF*^ and ultimately the transition probabilities **A** for the CHMM when using the total tree length *ℒ* as the hidden state. Similarly, the initial distribution can be obtained using equation (8).

#### 2.3.2 Emission Probabilities

Computing the emission probabilities closely follows the steps for the transition probabilities outlined in the previous section. However, we use the ancestral process with mutation *𝒜*^*θ*^ instead of the process with recombination *𝒜*^*ρ*^, and instead of one variable for time and two for tree length (*t, x, y*), we only need to use one variable for time and one for tree length (*t, x*) since we only consider emission at one locus. Before we can compute the emission probabilities, we first need to compute the joint probability of accumulating a certain tree length by *t* and *𝒜*^*θ*^ (*t*) occupying a certain state:

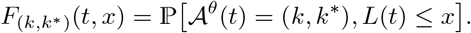

Here, *L*(*t*) is the accumulated tree length at this locus, defined similarly to equation (13) as

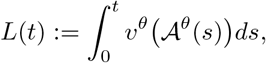

where *v*^*θ*^(*k, k*^∗^) = *k*. Similar to Section 2.3.1 and Miroshnikov and Steinrücken (2017), the vector of these probabilities can be obtained as the solution to the following system of PDEs

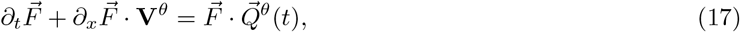

with boundary conditions

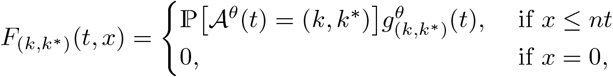

where 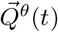 is the matrix of transition rates of the process *𝒜*^*θ*^ and the diagonal matrix

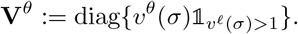

The solution to this system can be obtained using a similar approach to Miroshnikov and Steinrücken (2017), and we provide details in Section 1 in S1 Text. Similar to equation (9), we can then combine the probabilities for the absorbing states with the respective combinatorial factors to obtain the joint probability distribution of the tree length *ℒ* and the observed number of derived alleles as

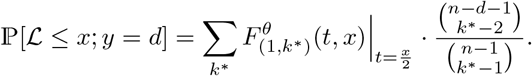

We can then again evaluate these probabilities at the discretization points *x*∈𝒮 = {*t*_0_, *t*_1_, …, *t*_*S*_} to obtain the entries of the matrix of cumulative probabilities **B**^*CDF*^, which can be substituted into equation (10) and (11) to obtain **B**^*PDF*^, and ultimately the emission probabilities **B** for the CHMM using *ℒ* as the hidden state, that is, the probabilities of observing a certain number of derived alleles, given the tree length.

### 2.4 Inferring Model Parameters

In this section we detail the procedure for inferring demographic model parameters using the HMM framework introduced in the previous section and introduce some extensions of the algorithm.

#### 2.4.1 Expectation-Maximization Algorithm

We use the Expectation-Maximization (EM) algorithm for HHMs to iteratively infer 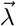, the parameters of the coalescent rate function *λ*(*t*) (and consequently the population size history *η*(*t*)). For *λ*(*t*), we use either a piecewise constant parametrization or a spline parametrization as described in more detail in Section 2.1 in S1 Text, and thus we infer a finite number of parameters. We choose the discretization for the hidden states independent from the population size history. We provide details in Section 3 in S1 Text. Briefly, for *T*_MRCA_, we use a discretization roughly equidistant on an exponential scale, and for *ℒ*, we choose a discretization such that the marginal distribution over the hidden states under a constant population size is approximately uniform.

We denote the parameters in the *k*-th iteration by 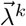. The initial parameters 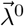 can be specified either by the user or by Watterson’s estimator (see Section 2.2 in S1 Text). For the *k*-th iteration of the E-step, we compute the initial (**Π**), transition (**A**), and emission (**B**) probabilities under the coalescent rate function given by 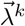. In the case of *T*_MRCA_ as the hidden state, we use a Dormand-Prince algorithm of order 8(5,3) (Dormand and Prince, 1980), to solve the respective ODEs. When using *ℒ* as the hidden state, we compute the probabilities by solving the associated PDEs using the scheme detailed in Section 1 in S1 Text, where we again use the Dormand-Prince method for the boundaries that require solving ODEs.

Using these probabilities, we then apply the Forward-Backward algorithm (Bishop, 2006, Ch. 13.2.2) to the observed genotype data 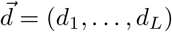, where *dℓ* is the number of derived alleles at locus *ℓ*. In Section 5 in S1 Text, we explain how we process input from `vcf`-files to obtain this vector of derived allele counts. The Forward-Backward algorithm yields the likelihood of the current demographic parameters, and its results can be used to compute 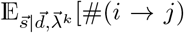, 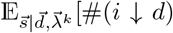, and 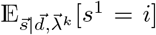, the number of expected transitions from state *i* to *j*, expected emissions of *d* derived alleles given state *i*, and the expected initial state, respectively, all conditional on the current parameters and the data. We use the scaled implementation (Bishop, 2006, Ch. 13.2.4) for numerical stability. Evaluating the Forward-Backward algorithm at each nucleotide site in the genome can become prohibitive. We thus detail two strategies to speed-up these computations in Section 4 in S1 Text: a locus-skipping method that compresses the computations between segregating sites, and a meta-locus method that groups segments of the genome into meta-loci to reduce the effective number of loci.

After each E-step we perform an M-step during which we update the values of 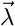 by numerically maximizing the objective function, defined as the expected log-likelihood of 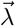 with respect to the conditional distribution of the hidden states given the data and the current parameter estimates 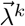,

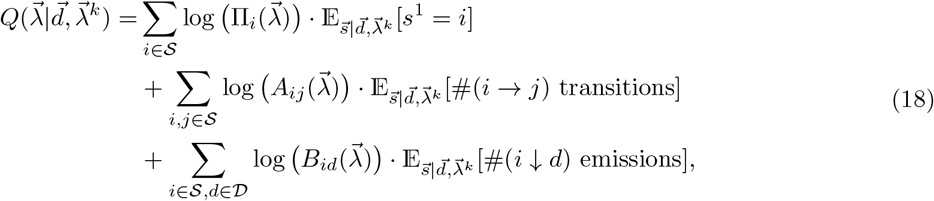

where we explicitly denote the initial, transition, and emission probabilities as functions of 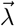 to stress that they are computed for the parameters that we optimize over. The parameters that maximize this function and thus yield the updated parameters for the next iteration are given by

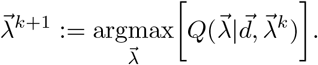

Since `CHIMP` evaluates the *Q* function numerically, we use the Nelder-Mead simplex optimization procedure for numerical optimization (Nelder and Mead, 1965). We observed some dependence of the results on the shape and orientation of the intial simplex for the Nelder-Mead procedure in each M-step, as the overall search direction can be biased by the orientation. To alleviate this bias, we initialize each M-step using the procedure described by Spendley et al. (1962), if the number of parameters to estimate is less than 5, and the procedure described by Gao and Han (2012) otherwise, as implemented in `scipy.optimize`. The former creates the initial simplex by adding and subtracting fixed values to the previous optimum along the coordinate axis, whereas the latter initializes the simplex by adding percentages of the previous values along the coordinate directions. While this distinction based on number of parameters seems unintuitive, we found that the performance differed substantially, and this approach performed best.

The optimization is performed in a search space of logarithmic coalescence rates, which is a uniquely robust space in which to perform optimization of coalescent rates (Parag and Pybus, 2019) and also has the benefit that the parameters for the coalescent rates are positive by design. In Section 2.3 in S1 Text, we provide details on different implementations to possibly regularize the population size function in the inference, however, no regularization was used for the simulation studies presented in Section 3. After finding the optimal coalescent rates using the EM algorithm, we invert and scale them to recover the estimates for the population size history *N* (*k*).

#### 2.4.2 Composite Likelihood

Our method takes as input genotype data of a sample of *n* haploids at *L* consecutive sites of the genome. However, in addition to applying the EM algorithm to all sampled haploids, we can also define a composite likelihood optimization scheme as follows. We can choose a set of subsets of the given haploids. These subsets can differ in size and can overlap each other, or be non-overlapping. In the E-step, for a certain subset of size *n*_*s*_ and given 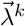, we can then compute 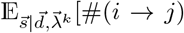 transitions], 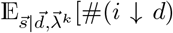 emissions], and 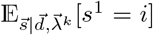 where *d ∈* {0, …, *n*_*s*_}, and compute the *Q* function in equation (18) based on the expected values for just this subset. We can then sum the *Q* functions across all subsets to obtain a composite function that is then maximized in the M-step. Repeating these EM-steps until convergence thus maximizes a composite likelihood, where the likelihoods of the subsets are multiplied.

This is procedure is useful in two ways. Firstly, subsets of different sizes are expected to have their *T*_MRCA_ at different times in the past, and the expected length of the trees will also differ. Thus, subsets of differing sizes potentially yield power to infer the size history in different periods. We explore this idea in Section 3.1. Secondly, the numerical procedures to compute the transition and emission probabilities are more computationally expensive for larger sample sizes. Thus, this composite likelihood scheme also allows us to more efficiently analyze large samples. The E-step scales linearly with the number of samples in each subset. Moreover, the computational time of the M-step depends only on *n*_*s*_, which is especially beneficial if the M-step is computationally more expensive (which we found to be the case when using *ℒ* as the hidden state). Thus, using this composite approach with smaller subsets, rather than analyzing the whole sample at once, decreases runtime substantially.

We also use this composite likelihood scheme to perform parameter inference using sequence data from different chromosomes simultaneously by aggregating the conditional expectations across chromosomes. In addition, this composite likelihood scheme is closely related to `MSMC2` (Wang et al., 2020), as the authors use all overlapping sub-groups of size *n*_*s*_ = 2 for the E-Step, and combine them in a similar way for the M-Step. In our software implementation, we allow the user to specify the subsets for this composite likelihood in two ways. The user can specify a list of sizes. for each of the given sizes, the complete sample is divided uniformly at random into non-overlapping subsets of the given size. The composite likelihood then multiplies across all subsets of a given size, and also multiplies across sizes. Alternatively, the user can specify an input file that explicitly lists the subsets (of potentially differing size) to be multiplied. The latter can be used to define overlapping subsets. We choose to not implement overlapping subsets as command line options, as the combinatorics can quickly become prohibitive.

## 3 Results

Before presenting and application of our method to population genomic data for humans from the 1000 Genomes project (Byrska-Bishop et al., 2021), we evaluate the accuracy of our method by performing a series of simulation studies on data generated under various demographic scenarios. We inferred the population size history from these simulated datasets using `CHIMP` with *T*_MRCA_ and *ℒ* as the hidden state under different composite likelihood schemes, and compared the results to inference using `MSMC2` (Wang et al., 2020, v2.1.2) and `Relate` (Speidel et al., 2019, v1.1.3). Note that we use `MSMC2` and `Relate` to infer the size history of a single population, but the methods can also be applied to samples from multiple populations to characterize population structure. For each study we used the specified model of the demographic history to simulate *m* = 16 replicates of data using `msprime` (Kelleher et al., 2016, v1.0.4), where each replicate consists of *n* = 200 haplotypes of length 200 Mbp. The per generation per site recombination and mutation rates we used were *r* = *μ* = 1.25 · 10^*−*8^, to mirror applications to human genetic data. We inferred the population size history for each of the replicates using the different methods and visualized the variability of the estimates across replicates. The scenarios presented here all require the inference of 15 or more parameters. If no prior information is available, a common strategy in the literature is to estimate a piecewise-constant size history with many changepoints, to be able to capture most relevant features of the true underlying history. We explore inference in a bottleneck and a piecewise growth scenario where we restrict to few given changepoints and a low number of parameters to be inferred in Section 7 in S1 Text.

We note here that the performance of `Relate` improved when we simulated and analyzed data with a human recombination map (see Section 8 in S1 Text). This is likely due to the fact that `Relate` benefits from *cold* spots (regions of low recombination rate) in the recombination map. However, the performance of `CHIMP` and `MSMC2` were not substantially adversely affected when they were run with the (inaccurate) assumption of a constant recombination rate. For this reason, we proceeded with a uniform recombination map in our simulation studies. The current implementation of `CHIMP` does require the user to specify a genome-wide recombination rate. Since the results of `CHIMP` were not affected much by varying recombination rates, we also expect the method to be resilient against misspecification of this parameter.

We first explore different composite likelihood schemes for `CHIMP`, and then compare `CHIMP` against the other methods in the different demographic scenarios. In each case, we either use either the full 200 haplotypes simulated, or use a subset of 10 haplotypes chosen uniformly at random. We use `CHIMP`-*𝒯* to indicate when *T*_MRCA_ is used as the hidden state, and `CHIMP`-*ℒ* when *ℒ* is used. Moreover, we use subscripts to indicate the (composite) likelihood scheme used. `CHIMP`-*𝒯*_10_ and `CHIMP`-*ℒ*_10_ indicate the use of nonoverlapping subsets of size 10, whereas `CHIMP`-*𝒯*_2,5,10_ and `CHIMP`-*𝒯* _2,5,10_ indicate the composite likelihood multiplying across all non-overlapping subsets of size 2, all non-overlapping subsets of size 5, and all nonoverlapping subsets of size 10. Thus, `CHIMP`-*𝒯*_10_ for a sample of size *n* = 10 is just the likelihood using *T*_MRCA_ as the hidden state, whereas `CHIMP`*𝒯*_10_ for a sample of size *n* = 200 is the composite likelihood multiplying across all non-overlapping subsets of size 10. Unless specified otherwise we use non-overlapping subsets.

The method `MSMC2` can analyze all pairs of haplotypes in the dataset, which we did for the samples of size *n* = 10, but was computationally prohibitive for samples of size *n* = 200. For the later case, we instead restricted `MSMC2` to analyze a subset of 50 non-overlapping pairs of haplotypes (100 haplotypes total) since memory requirements of the method became a limitation beyond this number. Moreover, in an effort to ensure a fair comparison between the methods, we ran all analyses for a piecewise constant parametrization of the population size history, and chose the same change points across the methods. The change points could be explicitly specified for `CHIMP` and `Relate`. For `MSMC2`, specifying change points was achieved by providing a time-segment pattern (as required by the method) that placed the change points as close to the desired ones as possible. This yielded a close match in most cases, with minor inaccuracies in more recent times and very ancient times. To intialize the iterative inference methods, we chose Watterson’s estimator for `CHIMP`, as detailed in Section 2.2 in S1 Text, and the default initialization for `MSMC2`.

To aid visualizing and summarizing the performance of a method in a specific setting, as well as comparing the results between methods, we also plot the mean absolute deviation from the true population size in generation *k*, or mean signed error, across the replicates

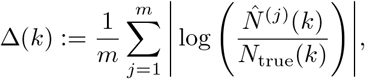

where 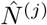 is the population size estimated in replicate *j*, and for each method, compute the integral of this quantity 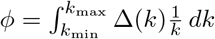 as a measure of discrepancy from the truth over the full history. Here *k*_min_ and *k*_max_ are the minimum and maximum of the respective discretizations with one logarithmic discretization step subtracted and added, respectively. Note that the factor 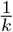 suppresses deviations in the distant past and transforms the integral to a regular integral of Δ on a log(*k*) timescale, which matches the visualization more closely.

### 3.1 Evaluating Composite Likelihood Approaches

A benchmark for population size inference that has been used in recent studies is a population size history that exhibits oscillations, referred to as *sawtooth* history (Schiffels and Durbin, 2014; Terhorst et al., 2017; Speidel et al., 2019; Sellinger et al., 2021). We will analyze a continuous version of this scenario in Section 3.3, but we were first interested in comparing the performance of the methods for a piecewise constant version so that the true population size history could in principle be exactly recovered using the different methods. To this end, we simulated data under a piecewise constant sawtooth history, were the population size oscillates between 50,000, 15,811, and 5,000 at 14 change points that are equidistant on a logarithmic scale between 57 and 448,806 generations before present. We simulated 16 replicates for this scenario.

We sampled *n* = 10 haplotypes uniformly at random from the simulated data and inferred the population size history using `MSMC2`, `CHIMP`-*𝒯*_2_ with overlapping subsets, and 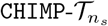 for *n*_*s*_ *∈* {2, 5, 10} with non-overlapping subsets. In addition, we performed the same comparisons with *ℒ* as the hidden state instead of *T*_MRCA_. The results are shown in Figure 5. We observe that `CHIMP`-*𝒯*_2_ with overlapping subsets performs almost identical to `MSMC2`, which is expected, since the two methods use essentially the same composite likelihood model. There are however striking differences in the very recent and very ancient times. We believe that these differences are due to the optimization scheme used in the M-Step, and both approaches don’t have much power in the respective time periods. The method `MSMC2` uses a Powell optimizer, whereas `CHIMP` uses a Nelder-Mead scheme. While developing our method and experimenting with different optimization schemes we noticed that Powell behaves erratically if it has little power, whereas Nelder-Mead does not change the initial value, which results in the observed patterns.

**Figure 5:**
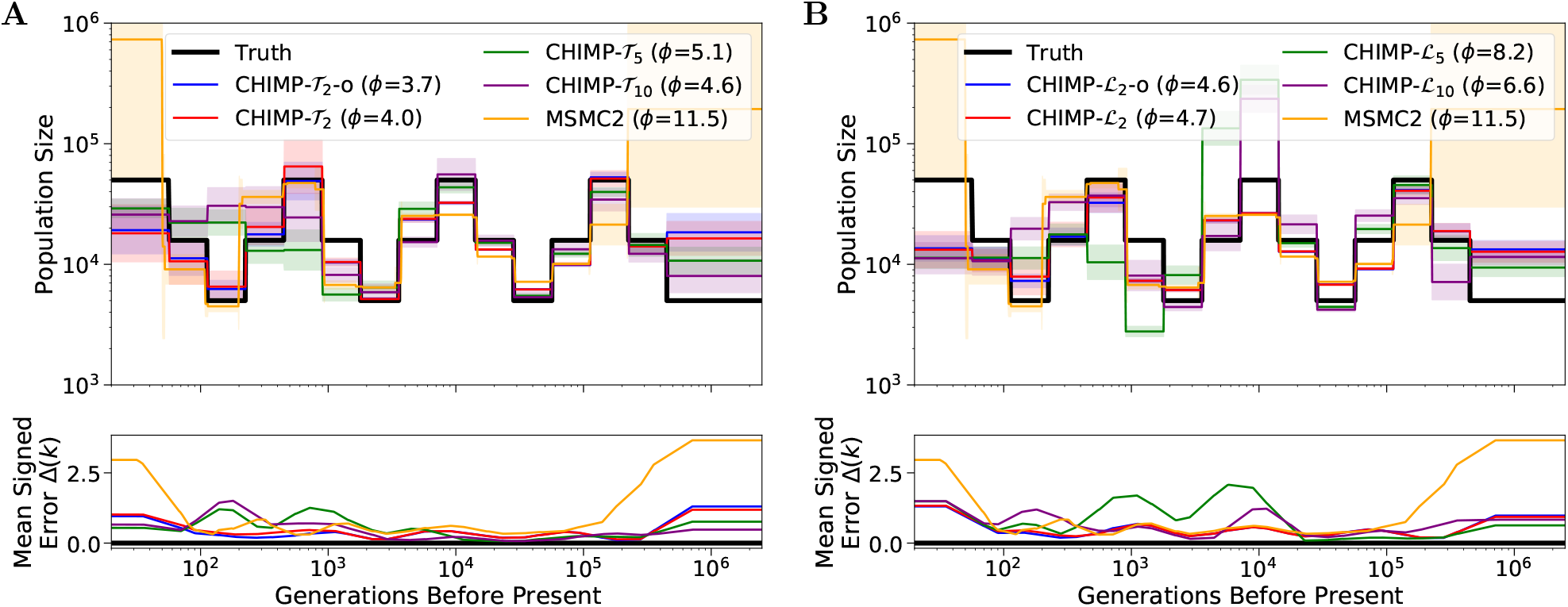
Results of inference in the piecewise sawtooth scenario from a sample of size *n* = 10 for different subset sizes using either *T*_MRCA_ (Panel **A**) or *ℒ* (Panel **B**) as the hidden state. We infer the population sizes in the intervals, fixing the change points to match the truth (shown in black). For `CHIMP`, we use non-overlapping subsets of sizes *n*_*s*_ = 2, 5, and 10. For *n*_*s*_ = 2, we also present overlapping subsets (-o). We present results obtained using `MSMC2` for comparison. Solid lines are averages over 16 replicates and the standard deviation is indicated by the shaded areas. Mean signed error Δ(*k*) is shown in bottom plot and has been smoothed using moving average for visualization purposes. The integral *ϕ* is indicated in the legend. Note that `MSMC2` groups epochs in the very distant past due to limits of the method interface.

Furthermore, note that `CHIMP`-*𝒯*_2_ performs very similar, regardless of whether overlapping or non-overlapping subsets are used. We conclude that using overlapping subsets adds little information, if the data is simulated from a panmictic population, as is the case here, but could differ more when analyzing real datasets. Nonetheless, we only use non-overlapping subsets in the remainder, since this reduces the runtime substantially. Lastly, note that for smaller subsets *n*_*s*_ = 2, the method performs better in the recent time, whereas larger subsets, the performance in the ancient time is better. Again, this might is expected, since samples of smaller size have a more recent *T*_MRCA_. However, the improvement in ancient times for larger subsets is not as pronounced as the improvement for small subsets in the recent times.

For `CHIMP`-*ℒ* we observe similar trends, but the overall performance is worse. Especially for *n*_*s*_ = 5 and 10, the population size is strongly overestimated around 10, 000 generation before present. This is likely due to the fact that we infer many demographic parameters (when compared to the inference in Section 7 in S1 Text) which results in a high dimensional inference problem with a likelihood surface that is more difficult to navigate and causes the method to converge to a local optimum. The fact that the direction of the bias replicates over different datasets suggests that the initial parameter choice and the details of the numerical optimization procedure (Nelder-Mead algorithm) affect the navigation to the local optima. Note that `CHIMP`-*ℒ*_2_ performs slightly worse than `CHIMP`-*𝒯*_2_, which use the identical composite likelihood model. This is because the former performs up to 15 EM-Steps, whereas the latter performs up to 25, the default parameters for our method. While this is suboptimal here, it did result in a better overall performance of the methods.

Motivated by these results, we further investigate possible composite likelihood schemes. Since *n*_*s*_ = 2 performs well in the recent past, whereas *n*_*s*_ = 10 perform better in the ancient past, we aimed to combine these approaches into a scheme that performs well across all times. We thus analyze the same datasets using `CHIMP`-*𝒯*_2,10_, `CHIMP`-*𝒯*_2,5,10_, `CHIMP`-*ℒ*_2,10_, and `CHIMP`-*ℒ*_2,5,10_, that is, in addition to multiplying the composite likelihoods for different subsets of the same size, we multiply them across sizes as well, and show the results in Figure 6. While these composite likelihood schemes do perform better then *n*_*s*_ = 2 in the ancient past, and better then *n*_*s*_ = 10 in the recent past, they do not fully retain the best performance of their components. Since `CHIMP`-*𝒯*_2,5,10_ performs best overall, we do include this scheme in the comparisons in the remainder. It is possible that the performance could be improved by weighing the different subsets differently in the composite likelihood, but we leave such exploration for future work.

**Figure 6:**
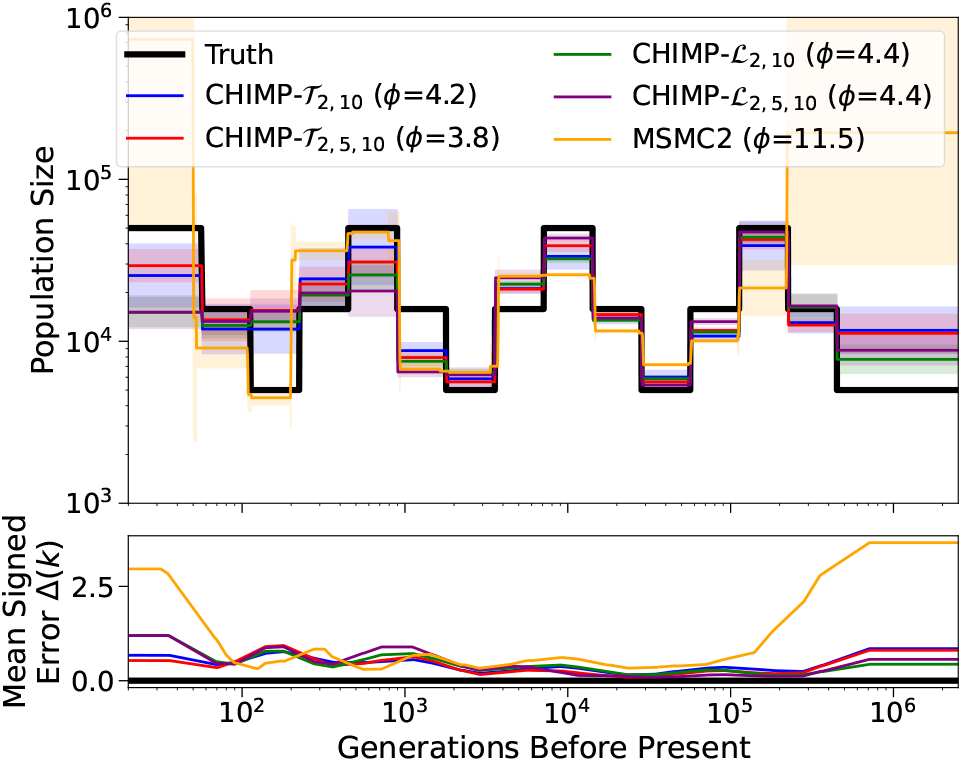
Results of inference in the piecewise sawtooth scenario from a sample of size *n* = 10 for the composite likelihood schemes `CHIMP`-*𝒯*_2,10_, `CHIMP`-*𝒯*_2,5,10_, `CHIMP`-*ℒ*_2,10_, and `CHIMP`-*ℒ*_2,5,10_. In these cases, the likelihood is multiplied across non-overlapping subset of the respective sizes, and multiplied across sizes. We present results obtained using `MSMC2` for comparison. We infer the population sizes in the intervals, fixing the change points to match the truth (shown in black). Solid lines are averages over 16 replicates and the standard deviation is indicated by the shaded areas. Mean signed error Δ(*k*) is shown in bottom plot and has been smoothed using moving average for visualization purposes. The integral *ϕ* is indicated in the legend. Note that `MSMC2` groups epochs in the very distant past due to limits of the method interface.

### 3.2 Compare Inference for Piecewise Constant Sawtooth Demography

In this section, we analyze the samples simulated under the piecewise constant sawtooth history using a uniformly sampled subset of size *n* = 10 and the full sample of size *n* = 200 with the methods `CHIMP`, `MSMC2`, and `Relate`. The results of this comparison are depicted in Figure 7. We observe that `CHIMP`-*𝒯*_2,5,10_ estimates the population sizes well across all times, except for some smoothing some recent times. `CHIMP`-*ℒ*_10_ estimates the size history accurately in the intervals 500 generations before present and further in the past, but also smooths the history in the very recent intervals. In general, `CHIMP`-_10_ behaves more erratically. It also does not infer the very recent times correctly, and is only correct for some of the intermediate intervals. The accuracy does not change substantially when using samples of different sizes.

**Figure 7:**
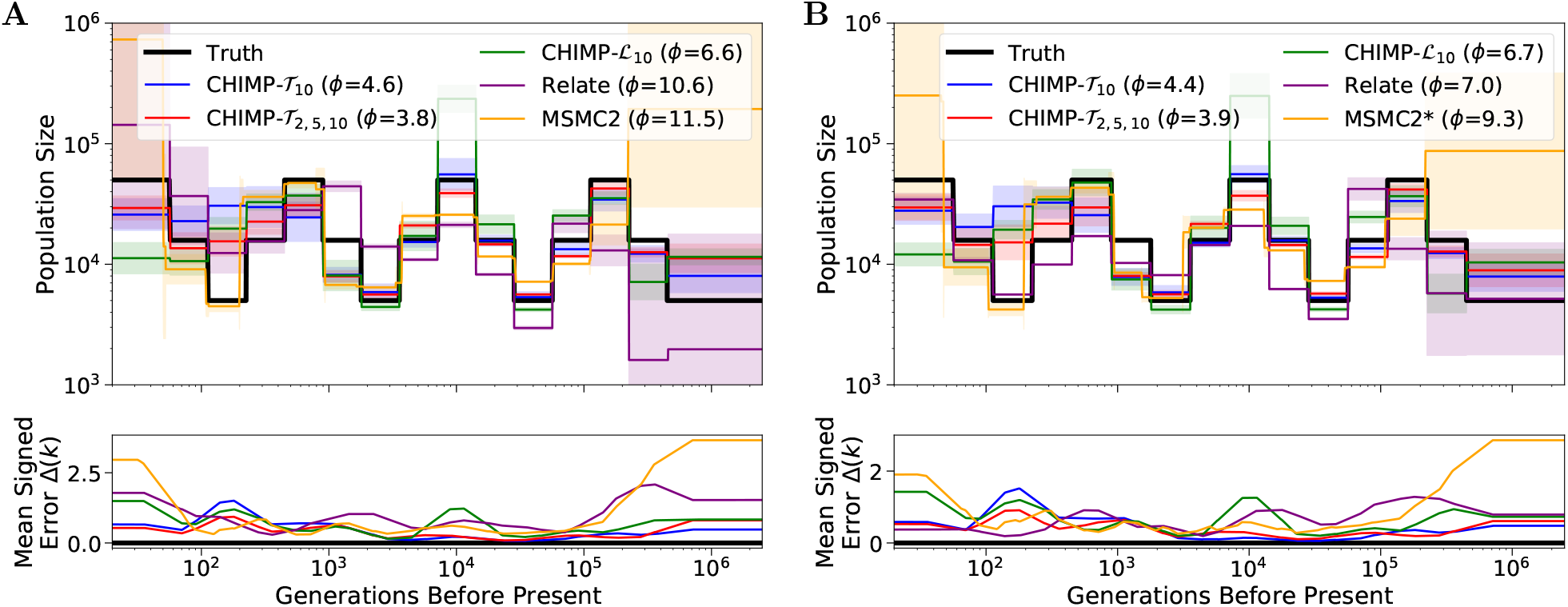
Results of inference in the piecewise sawtooth scenario for sample size 10 (Panel **A**) and 200 (Panel **B**). We compare the results using `CHIMP`, `MSMC2`, and `Relate` to infer the population sizes in the intervals, fixing the change points to match the truth (shown in black). Solid lines are averages over 16 replicates and the standard deviation is indicated by the shaded areas. Mean signed error Δ(*k*) is shown in bottom plot and has been smoothed using moving average for visualization purposes. The integral *ϕ* is indicated in the legend. Note that `MSMC2` groups epochs in the very distant past due to limits of the method interface. (*) For sample size 200, `MSMC2` was run on 50 non-overlapping pairs.

`MSMC2` shows accurate performance for intermediate times despite smoothing out some of the oscillations, but demonstrates high variability and a systematic upward bias below 100 generations and above 100,000 generations before present. Its accuracy does not change much between analyzing samples of different sizes. `Relate` has a high variability and upward bias for very recent times if the sample size is low, but the recent sizes are very accurately estimated when using a large sample size. The accuracy for intermediate and ancient times is not very high, and this performance is only slightly improved in intermediate times for larger sample sizes. Ultimately, between the different methods tested, each performs better in some timeframe or for specific sample sizes and worse for others. Measured in terms of integrated mean signed error *ϕ*, `CHIMP`-*𝒯*_2,5,10_ shows the overall best performance in this demographic scenario.

### 3.3 Inference for Continuously Varying Population Size History

In addition, we studied the performance of the inference methods on models of continuously varying population size history. Specifically, we considered the (continuous) *sawtooth* model implemented in `stdpopsim` by (Adrion et al., 2020, *ID=Zigzag 1S14*). In this model, the population size alternates between a maximum of 14,312 and a minimum of 1,431, with three maxima and three minima equidistant on a logarithmic scale between 33 and 34,133 generations before present. Note that the maxima and minima are roughly a fifth of the ones used by Schiffels and Durbin (2014) and Terhorst et al. (2017). We nonetheless decided to use this model here to investigate performance over a wider range of demographic scenarios in our simulation study. The second model we considered here is a bottleneck followed by exponential growth, a cartoon of an Out-Of-Africa population size history (Gutenkunst et al., 2009; Jouganous et al., 2017). In this model, the ancestral population size of 10,000 sharply drops to 2,000 at 4,000 generations before present. At 1,000 generations before present, the population size starts growing up to the present at an exponential rate of 0.25% per generation. Again, we simulated 16 replicates in each scenario with 200 haplotypes of length 200 Mbp and analyzed each replicate with each method on the full sample and on a subsample of size 10. We used the same discretization across methods for a better comparison, first specifying a minimum and maximum time and then choose 19 equidistant change points between these values (inclusive) on a logarithmic scale. The minimum and the maximum time were 40 and 40,000 for the sawtooth, and 200 and 20,000 for the bottleneck followed by growth scenarios, respectively.

The results in the sawtooth scenario are shown in Figure 8. We observe that for `CHIMP` and `MSMC2`, the accuracy does again not differ substantially between the different sample sizes. Again, the three versions of `CHIMP` smooth the population sizes earlier then 200 generations before present. Among these, `CHIMP`-*𝒯*_2,5,10_ follows the truth most closely, whereas `CHIMP`-*𝒯*_10_ and `CHIMP`-*ℒ*_10_ underestimate the very recent size 50 generations before present. The first peak, the second peak, and the ancestral population size are captured accurately by `CHIMP`-*𝒯*_2,5,10_, whereas `CHIMP`-*𝒯*_10_ and `CHIMP`-*ℒ*_10_ show some inaccuracy around the first peak. `MSMC2` captures both peaks, but slightly overestimates the ancestral size and substantially overestimates the recent sizes with a high degree of variability between replicates. For a small sample size, `Relate` overestimates recent sizes. It does infer two peaks, but the sizes and timing do not fully align with the truth. For a large sample size, `Relate` infers recent population sizes with high accuracy, but still underestimates the sizes of the two peaks. In terms of integrated mean signed error *ϕ* summarizing the overall accuracy, `CHIMP`-*𝒯* _2,5,10_ performs best.

**Figure 8:**
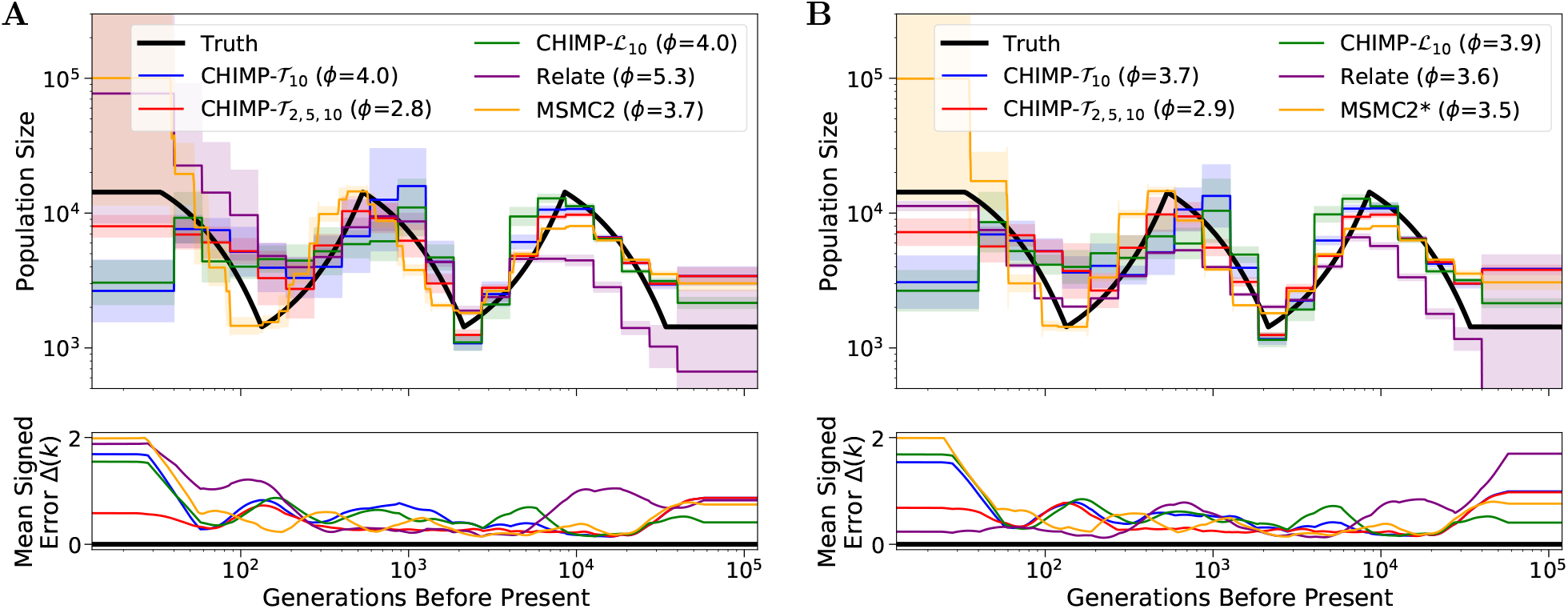
Results of inference in the continuous *sawtooth* scenario for sample size 10 (Panel **A**) and 200 (Panel **B**). We compare the results of `CHIMP`, `MSMC2`, and `Relate` using a piecewise constant population size history with 19 change points. Truth shown in black. Solid lines are averages over 16 replicates and shaded areas indicate standard deviation. Mean signed error Δ(*k*) is shown at bottom and has been smoothed using moving average for visualization purposes. The integral *ϕ* is indicated in the legend. (*) For sample size 200, `MSMC2` was run on 50 non-overlapping pairs.

Figure 9 shows the results in the scenario where a bottleneck is followed by exponential growth. In this scenario, all five methods capture the general trend of the population size history. Again, the performance of `CHIMP` and `MSMC2` does not differ substantially between sample sizes used in the analysis, however, `Relate` overestimates the recent sizes when using a small sample, but underestimates the history when using a large sample. `CHIMP`-*𝒯*_2,5,10_, `MSMC2`, and `Relate` smooth out the abrupt decline of the population size at the beginning of the bottleneck, whereas `CHIMP`-*𝒯*_10_ and `CHIMP`-*ℒ*_10_ do infer a sharper decline, but oscillate too much and do not infer the correct timing. We believe that some of the oscillations of `CHIMP`-*T*_10_ and `CHIMP`-*ℒ*_10_ are caused by the fact that the piecewise constant history is segmented into many pieces in a short time period, and the methods loose power. It is interesting to note that incorporating smaller subsets into the composite likelihood `CHIMP`-*𝒯*_10_ via `CHIMP`-*𝒯*_2,5,10_ does seem to recover some power. All methods but `Relate` infer the correct ancient sizes in the very distant past. In this scenario, the method `MSMC2` exhibits the best overall performance metric *ϕ*.

**Figure 9:**
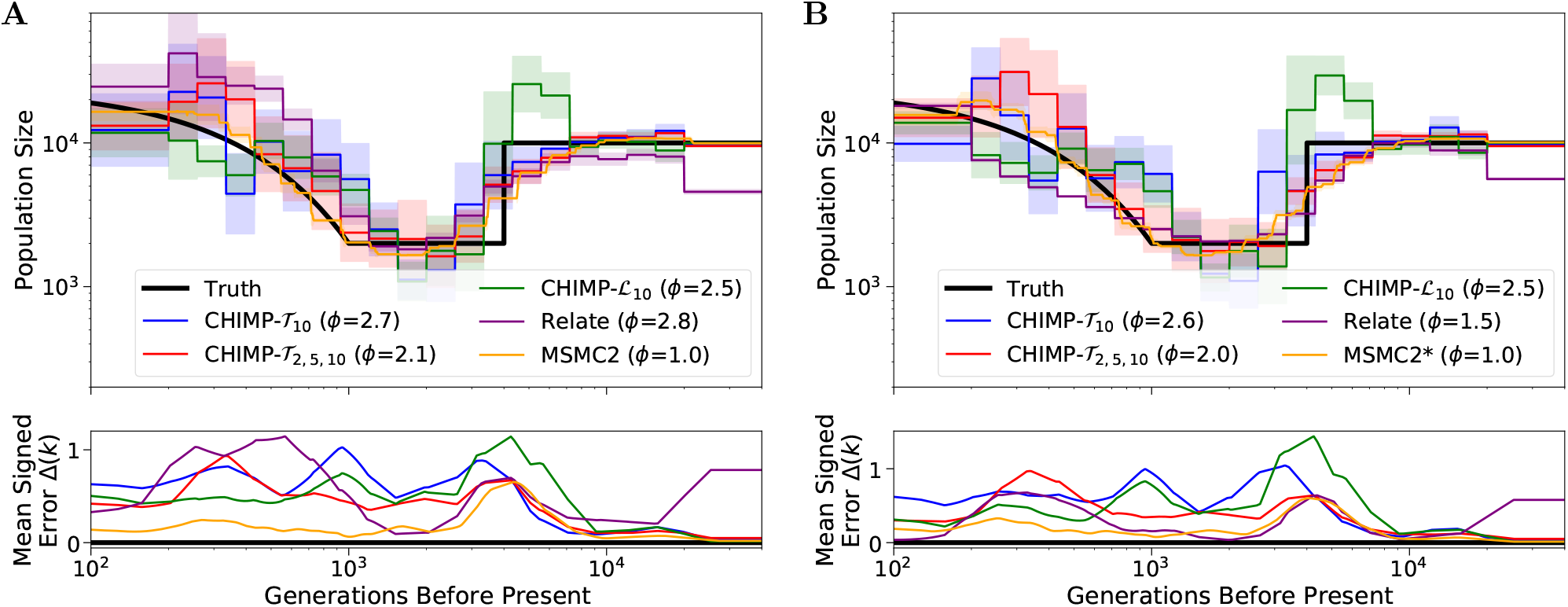
Results of inference in the bottleneck followed by growth scenario for sample size 10 (Panel **A**) and 200 (Panel **B**). We compare the inference of `CHIMP`, `MSMC2`, and `Relate` using a piecewise constant population size history with 19 change points. Truth shown in black. Solid lines are averages over 16 replicates and shaded areas indicate standard deviation. Mean signed error Δ(*k*) is shown in bottom plot and has been smoothed using moving average for visualization purposes, The integral *ϕ* is indicated in the legend. (*) For sample size 200, `MSMC2` was run on 50 non-overlapping pairs.

### 3.4 Computational Efficiency

In general, the runtime of the CHMM methods `CHIMP` and `MSMC2` scale linearly with number of loci *L*. In addition, the `CHIMP` methods scale linearly with the sample size, where we can use the composite likelihood framework introduced in Section 2.4.2 to reduce the effective sample size used in the computation of the transition and emission probabilities. If all pairs of samples are used in `MSMC2`, the method scales quadratically with the sample size, but when using non-overlapping pairs, like in our analysis of large samples, it scales linearly. `Relate` scales linearly with number of loci and quadratically with samples size. However, the method is implemented very efficiently and allows fast reconstruction of multi-locus genealogies for large sample sizes.

To exhibit the actual computational performance of the different methods in the simulation study, we list the average run-times in the different scenarios in Table 3. Since we ran `MSMC2` using 50 non-overlapping pairs of samples instead of the full 200, these are the times that we report. Additionally, runtimes for `MSMC2` are slightly inflated, as the number of CHMM states had to be increased to allow for the closest matching of demographic epochs. For all `CHIMP` methods, we observe a difference in performance between small and large samples, which is expected, since the number of subsets that are combined increases. We also note that the scenarios with few parameters to infer reported in Section 7 in S1 Text required less runtime than the more general scenarios. This is likely a result of fast convergence to an optimum when only a few parameters describe the model, whereas convergence is slower in a higher dimensional parameter space. The performance of `MSMC2` shows little variability across sample size and scenarios. Since for a sample of size *n* = 10, we analyze all 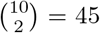 overlapping pairs, it is expected that the performance is similar to analyzing 50 non-overlapping pairs. The performance of `Relate` depends on the sample size, but shows little variability across scenarios, as expected, since the reconstruction of the genealogy is not strongly affected by the parameterization of the demographic model.

**Table 3:** Run-times in hours for the analysis of simulated data in the different scenarios, averaged over the respective 16 replicates in each case. The runtimes for `MSMC2` are slightly inflated, as the number of CHMM states had to be increased to allow for the closest matching of demographic epochs. (*) For the *n* = 200 scenarios, `MSMC2` was only run on 50 non-overlapping pairs of samples.

| | CHIMP-$\mathcal{T}_{10}$ | CHIMP-$\mathcal{T}_{2,5,10}$ | CHIMP-$\mathcal{L}$ | MSMC2* | Relate |
| --- | --- | --- | --- | --- | --- |
| Piecewise Sawtooth ($n = 10$, Fig. 7**A**) | 0.6 | 1.1 | 14.5 | 3.7 | 0.2 |
| Piecewise Sawtooth ($n = 200$, Fig. 7**B**) | 1.9 | 9.4 | 15.5 | 6.1 | 8.9 |
| Sawtooth ($n = 10$, Fig. 8**A**) | 1.0 | 1.7 | 19.8 | 4.1 | 0.1 |
| Sawtooth ($n = 200$, Fig. 8**B**) | 2.2 | 9.9 | 20.3 | 5.1 | 3.0 |
| Bottleneck + Growth ($n = 10$, Fig. 9**A**) | 1.0 | 1.3 | 19.3 | 5.2 | 0.1 |
| Bottleneck + Growth ($n = 200$, Fig. 9**B**) | 2.1 | 8.2 | 20.3 | 3.6 | 3.7 |

For sample size *n* = 10, `Relate` is the fastest method, but `CHIMP`-*𝒯*_10_ is only slightly slower. For *n* = 200, `CHIMP`-*𝒯*_10_ is fastest, but the runtimes of `CHIMP`-*𝒯*_2,5,10_, `MSMC2`, and `Relate` are on a similar order of magnitude. The method `CHIMP`-*ℒ*is substantially slower than the other methods. In these scenarios, with many parameters to be optimized, the EM algorithm requires many steps to converge, and each step requires evaluating PDEs to compute the transition and emission probabilities. This is computationally more expensive than, for example, evaluating ODEs as required for `CHIMP`-*𝒯*.

### 3.5 Analyzing Unphased and Pseudo-haploid Data

Our method `CHIMP` can be readily applied to unphased genomic data, and we will provide an explicit example in Section 3.7. It is therefore a promising method for applications where high quality phased genomes are not available, for example in human ancient DNA or for non-model organisms. In the simulation studies presented in this paper, we used simulated data, which is perfectly phased. However, the parameter inference performed using `CHIMP` only uses the number of derived alleles at each locus as input, and is therefore invariant to any phasing of the data. Thus, inference under the method will not be affected by phasing errors, and can even be performed on completely unphased data. `MSMC2`, when run on all possible of pairs of samples, requires phased data. If it is run on non-overlapping pairs of haplotypes where each pair is associated with a single individual, as we did here for large samples, it could be run on unphased data. However, such a scheme is not commonly used in the literature. Since `Relate` reconstructs multi-locus genealogies relating haplotypes, it cannot be applied to unphased data and will be adversely affected by phasing errors.

In addition to being able to analyze unphased data, our method can also take a form of pseudo-haploid data as input. Generating pseudo-haploid data is a strategy often applied to low-coverage sequencing data, where reliable diploid genotype calls are not feasible, and may introduce unwanted biases, for example in ancient human DNA studies (Barlow et al., 2020). In pseudo-haploid data, at each SNP, one sequencing read covering the respective SNP is chosen uniformly at random, and the allele on this read is then reported as the haploid genotype for the individual. We implemented an option for `CHIMP` that extends the CHMM to pseudo-haploid data. To analyze pseudo-haploid data for a sample of size *n*, we implement a two layered emission model. The CHMM is implemented using the *T*_MRCA_ for a sample of size 2*n* as the hidden state with the respective transition probability. At each locus, the emission in the first layer is then the number of derived alleles in a sample of size 2*n*. In the second layer, this sample is then down-sampled to a number of derived alleles in a sample of size *n* using hypergeometric probabilities.

We performed an additional simulation study, to demonstrate the inference from pseudo-haploid data using our method. To this end, we simulated 20 haplotypes of length 200 Mbp using the piecewise constant sawtooth demography (see Section 3.2). For each pair of haplotypes (diploid individual), at each locus, we selected one of the two alleles uniformly at random to obtain a dataset of 10 pseudo-haploid samples. We performed inference on the 10 pseudo-haplotypes of this data using `CHIMP`-*𝒯*_2,5,10_ with the pseudo-haploid option. We compared the results to those obtained using `MSMC2` and `Relate` for which we naively treated the data as if it were diploid data in order to perform demographic inference. The results are shown in Figure 10. Not surprisingly, `MSMC2` and `Relate` do not infer the correct population size history, with particularly large errors in recent times, whereas `CHIMP` retains an accuracy close that demonstrated for full diploid data. We note though that our method still relies on the information that no segregating sites are observed between the SNPs, which might not be available in low coverage sequencing data. However, we do believe that the capability to analyze pseudo-haploid data generated in this way presents an exciting avenue for future extensions.

**Figure 10:**
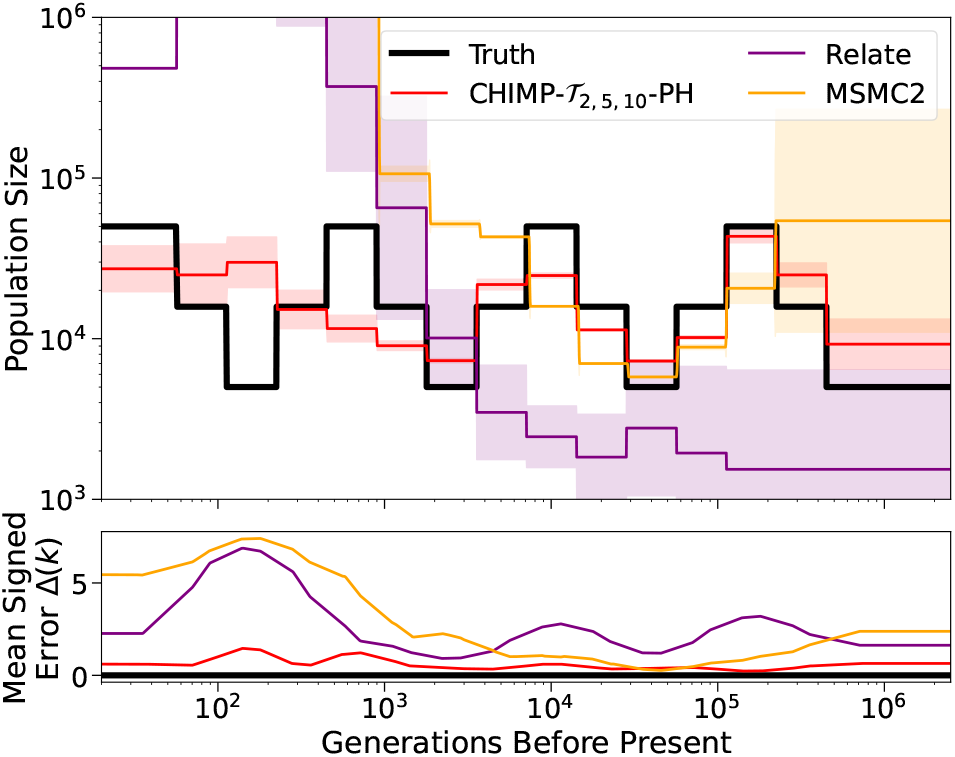
Results of inference using 10 pseudo-haploids simulated under the piecewise sawtooth demography. We compare the results of `CHIMP`-*𝒯*_2,5,10_ using the pseudo-haploid option, `MSMC2`, and `Relate`, fixing the change points to match the truth (shown in black). Solid lines are averages over 16 replicates and shaded area indicates standard deviation. Mean signed error Δ(*k*) is shown in bottom plot and has been smoothed using moving average for visualization purposes. The integral *ϕ* is indicated in the legend. Note that `Relate` entirely failed to estimate a population size in the most recent epochs, resulting in indeterminate values.

### 3.6 Summary of Simulation Study

Table 4 shows a summary of the performace and some features of the different methods that we compared in our simulation study. `CHIMP` and `Relate` can be applied to samples of arbitrary sizes, whereas `MSMC2` is limited in this regard. Furthermore, `CHIMP` can be applied to unphased data, and in limited capacity to pseudo-haploid data, but `Relate` requires phased data. `MSMC2` can be applied to unphased data, if the appropriate pairs of haplotypes are chosen for the analysis. `CHIMP`-*𝒯*_10_ and `Relate` can be used to analyze large samples quickly. `CHIMP`-*𝒯*_10_ is comparable, whereas `CHIMP`-*ℒ* was very slow. The runtime of `MSMC2` was comparable with `CHIMP`-*𝒯*_10_, `CHIMP`-*𝒯* _,5,10_, and `Relate`, but the method could not be run on the full sample of size *n* = 200. Moreover, `CHIMP` and `Relate` are very flexible in terms of user-specification of the demographic model, whereas `MSMC2` limits the user to choose an appropriate time-segment string.

**Table 4:** Summary of the performance and features of the different methods compared in our simulation study. The ranges are given in generations before present.

| | Gener. bp | CHIMP-$\mathcal{T}_{10}$ | CHIMP-$\mathcal{T}_{2,5,10}$ | CHIMP-$\mathcal{L}$ | MSMC2 | Relate |
| --- | --- | --- | --- | --- | --- | --- |
| Sample size |  | Large | Large | Large | Limited | Large |
| Parametrization |  | Flexible | Flexible | Flexible | Limited | Flexible |
| Accuracy | Few param. | High | High | High | High | Mixed |
| | $k < 50$ | Low | Mixed | Low | Low | High ($n \gg 1$) |
| | $50 < k < 500$ | Low | Mixed | Low | High | High ($n \gg 1$) |
| | $500 < k < 10^5$ | High | High | Mixed | High | Mixed |
| | $k > 10^5$ | High | High | Mixed | Low | Low |
| Runtime | | Fast | Fast | Slow | Fast (lim. $n$) | Fast |
| Unphased data |  | Yes | Yes | Yes | Possible | No |
| Pseudo-haploid |  | Limited | Limited | Limited | No | No |

The inference using `CHIMP` and `MSMC2` showed very high accuracy when analyzing scenarios with a limited number of demographic parameters. In contrast, `Relate` did not perform well in this case. When performing inference under a flexible piecewise constant parametrization, `CHIMP`-*T*_10_ has limited accuracy in recent and intermediate times, `CHIMP`-*𝒯*_2,5,10_ recovers some accuracy here. `CHIMP`-*ℒ* did perform worse in intermediate times, but all `CHIMP` methods perform well in ancient times. `MSMC2` did not infer the population size history well in recent and ancient times, but showed the best performance among the methods tested here in intermediate times, however, `CHIMP`-*𝒯*_2,5,10_ is a close second. `Relate` inferred recent population sizes well, if a large sample was used, but the performance was less accurate for small samples and intermediate to ancient times. We note that the exact time-frames depend on the baseline effective population sizes. The scenarios that we investigated here ranged from *N*_*e*_ ≈ 4, 000 to 12, 000.

Interestingly, the accuracy of `CHIMP` and `MSMC2` did not increase substantially when the methods were applied to samples of larger sizes. This is likely due to the fact that the application of the methods to larger samples is achieved in composite likelihood schemes that effectively only use smaller subsets, but might also indicate that in scenarios of truly panmictic populations, much of the population size history can be learned from just a few individuals. This is perhaps best exemplified by the immense success of PSMC (Li and Durbin, 2011), which extracts surprising amounts of information from just two haploid sequences of a single individual. `Relate` appears to require a certain minimal sample size to exhibit good performance, but demonstrates that for accurate inference of very recent population sizes, it is indeed necessary to sample many haplotypes.

In summary, `CHIMP`-*T*_10_ and `CHIMP`-*T*_2,5,10_ perform comparably to the other methods tested here in most scenarios when inferring sizes beyond 500 generations before present, and runs quickly on large datasets. `CHIMP`-*T*_2,5,10_ even recovers some accuracy below 500 generations before present. An advantage is the fact that these methods can be applied to unphased and, in limited capacity, pseudo-haploid data; thus they offer a useful alternative to other existing methods, especially in situations where high quality data is not available. Overall, the inference accuracy of `CHIMP`-*ℒ*_10_ was mediocre and the runtime was very poor. We thus do not recommend this approach for inference of populations size histories, unless improvements can be made in terms of efficiently computing the probabilities required for the CHMM and navigating the high dimensional optimization problem. `MSMC2` showed very high inference accuracy for intermediate times and thus proves to be an effective method if the sample size is not too large. `Relate` is fast and can be applied to large samples. The inference accuracy is good with sufficient samples, especially in recent times, but suffers if the recombination map does not have *cold* spots.

### 3.7 Inferring population Size History from Unphased Human Data

To demonstrate that our method can be readily applied to population genomic datasets, we analyzed sub-samples of the 1000 Genomes dataset recently re-sequenced to high coverage by Byrska-Bishop et al. (2021). In principle, this data has been computationally phased, but we did not use the phase information, to further demonstrate this feature of our method. Specifically, we downloaded the dataset in `vcf`-format from the server provided by the authors (see also `Data Availability`), and extracted genomic data for chromosome 1 of the individuals in the population groups `LWK` (African, 99 individuals), `JPT` (Asian, 104 individuals) and `FIN` (European, 99 individuals). Note that we did download the phased version of the data, but “removed” this information by switching each genotype uniformly at random. We decided on these unusual processing steps, since the phased data was more readily available for download without extensive preprocessing. We then inferred the population size history for each of these population groups using `CHIMP`, specifically the composite likelihood scheme `CHIMP`-*𝒯*_2,10_. We used the default parametrization of `CHIMP`, except for specifying the regularization coefficient *c*_12_ = 10^*−*5^, see Section 2.3 in S1 Text. We used a per generation mutation rate of *μ* = 1.25 · 10^*−*8^ and for the per locus per generation recombination rate we used the same value *r* = 1.25 · 10^*−*8^. The estimated population size histories are shown in Figure 11, were we used a generation time of 26.9, to convert generations into years as estimated by Wang et al. (2021).

**Figure 11:**
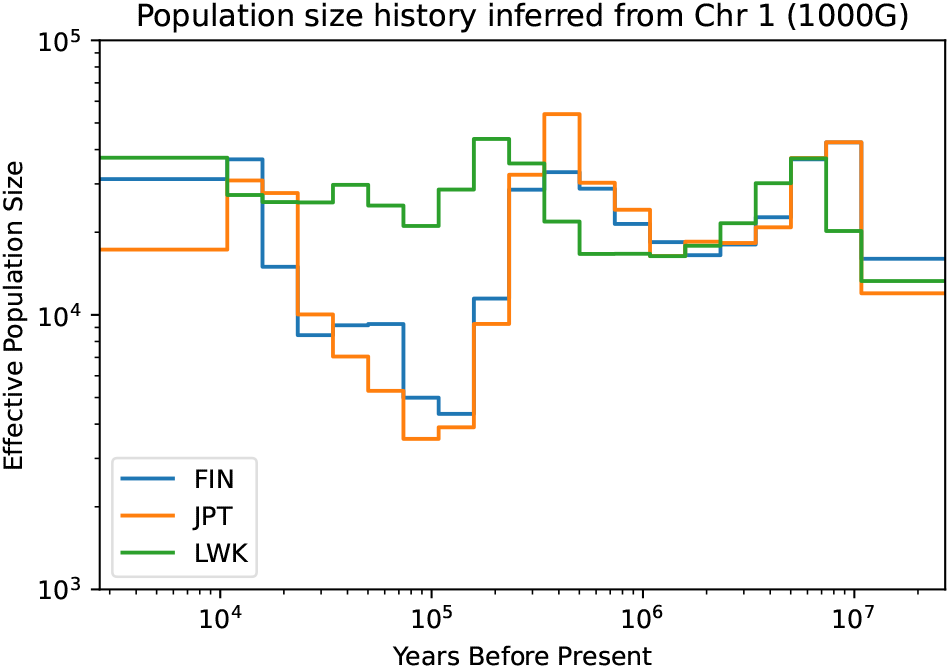
Effective population sizes estimated for the population groups `LWK`, `JPT`, and `FIN` from the 1000 Genomes dataset using `CHIMP`-*𝒯*_2,10_. The populations show a similar history up to approximately 200,000 years ago, when they start to diverge. The Non-African population exhibit the well characterized Out-Of-Africa bottleneck, with subsequent expansion in the recent past.

We observe that all three populations exhibit the a similar population history up to 200,000 years before present. Following this period, the population histories start to diverge. The history of the African population (`LWK`) stays at around the same level, whereas the population histories of the Non-African populations (`JPT` and `FIN`) undergo a severe bottleneck. Towards the more recent past, the population sizes increase again. This example shows that our method is able to recover well established features of the population size history of modern humans, like the Out-Of-Africa bottleneck, and subsequent exponential growth in Non-African populations, see for example Speidel et al. (2019).

## 4 Discussion

Here, we presented our novel flexible Coalescent HMM method `CHIMP` to perform inference of past population sizes in a single population. The method uses either the *T*_*MRCA*_ or the total branch length *ℒ* of the local genealogies as the underlying hidden state in this HMM framework. We detailed systems of differential equations derived from the ancestral process that can be used to compute the respective transition and emission probabilities. These differential equations can be computationally intensive to solve, particularly for *ℒ*, but we present solution schemes that exploit a combination of approximations and exact equations to obtain solutions. We also combine CHMMs for differently sized subsets of the sampled haplotypes to combine power in different time periods and speed up computation. Furthermore, the framework presented here can be seen as a generalization of most previous CHMM methods, in that it can readily be modified to use pairwise coalescent times (`PSMC` and `MSMC2`, Li and Durbin, 2011; Wang et al., 2020), first-coalescent times (`MSMC`, Schiffels and Durbin, 2014), as well as the coalescent time of a distinguished pair (`SMC++`, Terhorst et al., 2017) as the hidden state.

We applied `CHIMP` for demographic inference from simulated data in a variety of scenarios and compared the results to other state-of-the-art methods, specifically `MSMC2` and `Relate`. While `CHIMP`-*ℒ* is intriguing from a theoretical perspective, it currently does not seem suitable for demographic inference since inference is slow and less accurate than the other methods, although more efficient approximations may ameliorate this issue. Despite the long runtimes, `CHIMP`-*ℒ* is also outperformed by `CHIMP`-*𝒯* in most scenarios, albeit not substantially in some. We believe that this could be due to the fact that under the infinite sites model, the number of segregating sites is a sufficient statistic for the length of the tree, but not for the *T*_MRCA_. Our CHMM uses the number of derived alleles as the emission, and we believe that this provides more information about the underlying *T*_MRCA_ than about the length of the tree, thus resulting in better accuracy for `CHIMP`-*𝒯*.

We observed that `CHIMP`-*𝒯*_10_ performs comparably to other methods in most scenarios for time-frames more than 500 generations before present and outperformed them for very ancient times beyond 100,000 generations before present. Our composite likelihood approach `CHIMP`-*𝒯*_2,5,10_ performed as well for times greater than 500 generations before present, and shows adequate performance in earlier times. The runtimes of `CHIMP`-*𝒯* are similar to those of the other methods in the tests we performed, and it scales very well to large samples. `CHIMP` can also be run on unphased data and certain pseudo-haploid datasets, whereas `Relate` requires phased data, and `MSMC2` can only be run in a limited capacity without phased data. We believe, that this makes `CHIMP`-*𝒯* a flexible alternative to other methods when analyzing large data sets, especially in scenarios where high quality assessment of haplotype phase is not available, which includes non-model systems were reliable reference panels are not available. Other interesting approaches in this context include the recent extension of `Relate` by Speidel et al. (2021) that can estimate population size history for low quality ancient human DNA by borrowing power from a known genealogy of a high quality panel of related individuals.

We note that, when performing the analyses of simulated data presented here and designing comparisons between the methods that can be deemed fair, the flexibility of the user-interface and the heuristics to choose an a-priori discretization of the population size history for inference impacts the applicability and performance of the different methods. We showcase this with an application of the methods to the bottleneck followed by growth scenario in Section 6 in S1 Text, where we ran all methods using their default parameters. In Section 6 in S1 Text, we also detail the heuristic we implemented in `CHIMP` for determining a default parameterization. We found it to be robust in the scenarios that we considered and believe that it performs well in general. However, exploring optimal ways of parameterizing models with no prior information about the demographic history is an important area in which further study is needed. In this context, parameter free approaches, see for example work by Ki and Terhorst (2020), present interesting alternatives.

For practitioners, we advise applying a composite method that combines power like `CHIMP`-*𝒯*_2,5,10_ or `CHIMP`-*𝒯*_2,10_ to get good estimates for all time periods. If the more recent time is of interest, we would advise to complement such an analysis with `CHIMP`-*𝒯*_2_ (or `MSMC2`), and potentially `Relate`, if the data is of high quality and the sample size is large enough. The results of the simulation study presented here are obtained under certain assumptions about the underlying parameters, which we motivated by applications to humans. However, for organisms with different mutation or recombination rates, and different diversity levels, the exact details and power for inference at certain times in the past might differ. This will also be affected by the exact parametrization of the inference method. We thus stress that an analysis of empirical data using our method, or any other method for that matter, should be supplemented by simulation studies to establish the right parametrization and understand the behavior of the method in the respective scenarios.

We note that the assumption of only one recombination event per pair of adjacent nucleotide sites and one mutation event per nucleotide site are more likely to be violated with increasing sample size. From a practical perspective, we do not think that this is a major concern, since due to our composite likelihood scheme, the maximal sample size our method uses internally in the examples presented here is n=10. From a theoretical perspective, when assuming only one mutation event, despite several mutation events occurring on the respective tree, our method would tend to infer shorter trees, thus likely incurring a bias towards more recent coalescence, and underestimating population sizes. For parameters appropriate in human populations, these events are rare, and will likely not result in noticeable bias, but in organisms with higher mutation rates, this approximation has to be re-evaluated. Similarly, assuming only one recombination event would make correlation among the hidden states stronger, likely increasing the variance of the estimates, but potentially not introducing systematic bias.

The modeling framework developed here has potential applications beyond inferring the population size history of a single population. The differential equations that we presented to compute the requisite probabilities for the CHMM were derived from the ancestral process in a panmictic population, but the approach can be extended to structured populations by augmenting the state space, enabling inference of migration rates and divergence times, and characterizing admixture between populations. Moreover, samples at different points in time (e.g. ancient samples), can be readily incorporated into the ancestral process and thus the inference framework as well. In addition, using the model and possible extensions presented here to characterize the posterior distribution of the local genealogies has many interesting applications. Different forms of selection impact the local genealogies around beneficial alleles, and thus the posterior distribution of the *T*_*MRCA*_ or the total branch length *ℒ* can help identifying and characterizing adaptive genetic variation (Stern et al., 2019; Speidel et al., 2019).

## Data Availability

Our software `CHIMP` (**C**HMM **H**istory-**I**nference **M**L **P**rocedure) is available for download at https://www.github.com/steinrue/chimp. Instruction on how to run the software and scripts to recreate the simulation study presented here can be found at this url as well. For the analysis of the 1000 Genome dataset, we downloaded the genotype data from http://ftp.1000genomes.ebi.ac.uk/vol1/ftp/data_collections/1000G_2504_high_coverage/working/20201028_3202_phased/, the reference sequences from ftp://ftp.ncbi.nlm.nih.gov/genomes/all/GCA/000/001/405/GCA_000001405.15_GRCh38/seqs_for_alignment_pipelines.ucsc_ids/, and the ancestral alleles from ftp://ftp.ensembl.org/pub/release-104/fasta/ancestral_alleles/.

## Acknowledgements

We thank the Steinrücken, Novembre and Berg lab for many inputs and helpful feedback. We also thank Maryn Carlson and Arjun Biddanda for feedback on the manuscript. In addition, we would like to thank Margarita Orlova for assistance in performing the simulation studies.

## Supporting Information

### S1 Text. Supplementary Text for Robust Inference of Population Size Histories from Genomic Sequencing Data

This supplementary text contains additionals details on the derivation, on the implementation, and additional analyses.

