## Supplementary material for "Robust Inference of Population Size Histories from Genomic Sequencing Data": S1 Text

### S1 Text: Supplementary Text for Robust Inference of Population Size Histories from Genomic Sequencing Data

#### 1 PDE Solution Scheme

To numerically compute the requisite solutions to the PDEs (14) and (17) in the main text, we implement the solution scheme outlined by Miroshnikov and Steinrücken (2017). We will give a brief overview of the scheme here, but refer the reader to the original publication for more details. Moreover, we will focus on the solution scheme for the joint distribution of the tree length $F_\sigma(t, x, y)$ in equation (14) in the main text here, but the same scheme can be used for the emission distribution in equation (17) in the main text. In this scheme, $F_\sigma(t, x, y)$ is computed for each state sequentially, beginning with the initial state $(n, 0, 0, 0)$. Because we explicitly keep track of the number of recombination events, we can ensure that each transition of the associated Markov chain either increases the number of recombination events or decreases some number of lineages. The states are thus ordered, and following this order in the evaluation, we can ensure that when computing $F_\sigma(t, x, y)$, all preceding $F_{\sigma'}(t, x, y)$ that are necessary for the respective computations have already been computed. In other words, since there are no loops in the probability flow through states, we can solve them sequentially following the natural ordering.

To compute the solution for a specific $F_\sigma(t, x, y)$, we favor a bespoke implementation of the PDE solutions as opposed to using a black box numerical PDE solver, since the actual solutions for each state are computed using the Method of Characteristics. This allows us to take advantage of partially analytic results which can reduce numerical errors and increase efficiency. Moreover, in terms of efficiency, we believe solving the states sequentially is faster and more accurate than attempting to simultaneously solve for all states using a general purpose method.

For each state $\sigma = (k_{ab}, k_a, k_b, r)$ we then define a three dimensional grid of points for the function $F_\sigma(t, x, y)$, with $0 \leq t \leq \frac{t_{S-1}}{2}$, $v^a(\sigma) \cdot t \leq x \leq n \cdot t$, and $v^b(\sigma) \cdot t \leq x \leq n \cdot t$. The grid consists of slices in the $x$-$y$ plane that are spaced across $t$ according to a specified discretization $D := \left\{t_0 = 0, \ldots, t_K = \frac{t_{S-1}}{2}\right\}$ into $K$ values. Each successive grid along $t$ is defined by propagating along the direction of the characteristic $(x_0 + v^a(\sigma)\tau, y_0 + v^b(\sigma))$ from the previous grid, with a new column and row for $x = n \cdot t$ and $y = n \cdot t$ on the outer boundaries. Note that the first grid consists of the point $(0, 0, 0)$ only. The next step extends the characteristic out of this point and adds the correct values at the boundaries, and thus the initalization of the scheme is well defined.

To ultimately fill all grids with the correct values, we start at each grid point along the boundary and fill in values for $F_\sigma$ by integrating along the characteristic direction. This fills in lines of points in the interior of the volume, and once all boundary points are processed, all interior points will be computed. For $\sigma = (k_{ab}, k_a, k_b, r)$, take a point $(t_0, x_0, y_0)$ on the boundary, that is, $t_0 \in D$ (the discretization grid), and either $x_0 = n \cdot t_i$ or $y_0 = n \cdot t_i$. Moreover, let $F_\sigma(t_0, x_0, y_0) =: F_{\sigma 0}$. Then, for $(t_0 + \tau) \in D$, the values on the grids along the characteristic can be computed according to:

$$
\begin{aligned}
F_\sigma\big(t_0+\tau, x_0+v^a(\sigma)\tau, y_0+v^b(\sigma)\tau\big) &= e^{-H_k(\tau)}\Big(F_{\sigma 0}+\int_0^\tau g_\sigma(\alpha)e^{H_k(\alpha)}d\alpha\Big)\\
\text{where } g_\sigma(\tau) &:= \sum_{\sigma'\to\sigma} F_{\sigma'}\big(t_0+\tau, x_0+v^a(\sigma)\tau, y_0+v^b(\sigma)\tau\big)Q^\rho_{\sigma',\sigma}(t)(t_0+\tau),\\
H_\sigma(\tau) &:= \int_0^\tau q_\sigma(\alpha)d\alpha,\\
q_\sigma(\tau) &:= -Q^\rho_{\sigma,\sigma}(t_0+\tau), \text{ and}\\
\sigma' \to \sigma & \text{ is the set of states } \sigma' \text{ that precede } \sigma.
\end{aligned}
$$

This scheme implementing the Methods of Characteristic propagates the boundary values along the characteristics. While the integral for $H_\sigma(\tau)$ is computed exactly (since we can analytically integrate $\lambda(t)$ for spline and piecewise constant representations) the integral in the expression for $F_\sigma$ is computed by approximating the integrand as piecewise linear, with the pieces bridging points of the grid for $\sigma$ (along a characteristic). This allows us to restrict the evaluations of $g_\sigma$, $H_\sigma$, and $q_\sigma$ to points on the $\sigma$-grid.

As seen in equation (14) in the main text, the values along the boundaries are the solution to another set of PDEs, defined in equation (15) in the main text, albeit in 2 dimensions rather than 3. The boundary value PDEs can be solved using a similar scheme with the dimension reduced, following one dimensional characteristics (along which these boundary points lie). The boundary values for these secondary PDEs are $F_\sigma(t, x = y = n \cdot t) = g^\rho_\sigma(t)$ and can be computed by solving a set of ODEs using a standard numerical ODE-solver (as was done for $T_{MRCA}$).

We note that in solving for each state, the solutions for all states immediately upstream (states preceding $\sigma$ in the ordering) are required. The grid for each state is based on the same discretization $D$ of time-steps. However, the $x$ and $y$ values along the grids do depend on the characteristic direction which can differ between different states, and thus, for a given $t$, the $x$-$y$ grid for a state $\sigma$ may be different than the grid used for a preceding state $\sigma'$. Thus, in order to compute $g_\sigma(\tau)$, we obtain the values for $F_{\sigma'}$ on the $x$-$y$ grid for the state $\sigma$ by linearly interpolating the values from the grid of $\sigma'$. We also tested interpolation using a cubic spline, but it resulted in only marginal improvements in accuracy at the cost of significant computational time and potential numerical instabilities due to oscillations.

We furthermore improve efficiency by imposing the following symmetries and constraints explicitly in our solution scheme, which follow from symmetries in the ancestral process with recombination $\mathcal{A}^\rho(t)$:

$$
\begin{aligned}
F_{(k,r,r,r)}(t,x,y) &= F_{(k,r,r,r)}(t, \min(x,y), \min(x,y))\\
F_{(k,k',k',r)}(t,x,y) &= F_{k,k',k',r}(t,y,x)\\
F_{(k,1,0,1)}(t,x,y) &= F_{k,0,1,1}(t,y,x),
\end{aligned}
$$

for all $k \in \{1,\dots,n\}$ and $k', r \in \{0,1\}$. After computing $F_\sigma(t,x,y)$ for all $\sigma$ and all points on the grids, we can substitute the numerical values into equation (16) in the main text to compute the requisite probabilities for the transitions of our CHMM. Emission probabilities are computed using an analagous scheme for the PDE given in equation (17) in the main text.

### 2 Population Size Parameters

#### 2.1 Parameterization of $\eta(t)$

While the population size history $\eta(t)$ is in general a non-singular positive-valued function, in order to perform efficient inference we restrict $\eta(t)$ to be parameterized by a finite number of parameters in two different ways.

Our default option is to represent $\eta(t)$ as a piecewise constant function, where the number of pieces can be specified by the user and are uniform in $\log(t)$ space between specified bounds. In practice, it is more convenient to directly work with $\vec{\lambda}$, defined to be the piecewise values for the coalescence rate $\lambda(t)$, and transform these back into the population size at the end.

We also include an option to perform the inference using a cubic-spline representation of the population size history, similar to that offered by `SMC++` Terhorst et al. (2017). More specifically we represent $\lambda(t)$ as a cubic spline, and the population size history is proportional to the inverse of this. The number of nodes can be specified, and we place them equidistantly on a $\log(t)$ scale.

#### 2.2 Initialization of Population Size History

We initialize the EM algorithm to infer the population size history such that $N(k) = \hat{N}$ is constant over time. For this constant value $\hat{N}$, we use the effective population size inferred from the data using Watterson's estimator of $4\hat{N}\mu$, fixing the mutation probability $\mu$ to a value appropriate for the data analyzed. We also use the value for $\hat{N}$ to guide choosing a partition for the CHMM states (Appendix 3) and in computing regularizing coefficients (Appendix 2.3).

#### 2.3 Regularization of Inference Procedure

For demographic inference problems, we might have some prior notion of the shape of the population size history function. For different applications we may have different tolerances for sharp changes in population size, or may expect varying strengths of deviations from flat histories.

Including regularization parameters helps to suitably constrain the inferred population size histories. For a given choice of population size history parameters that define a coalescence rate function $\lambda(t)$, we define the following four regularization quantities, where $\hat{\lambda}$ is the coalescent rate corresponding to Watterson's estimate for the effective population size in a constant model, computed from the data:

$$
\begin{aligned}
R_{02}(\lambda(t)) &= \int \left(\lambda(t) - \hat{\lambda}\right)^2 dt \\
R_{11}(\lambda(t)) &= \int |\lambda'(t)| dt \\
R_{12}(\lambda(t)) &= \int \left(\lambda'(t)\right)^2 dt \\
R_{22}(\lambda(t)) &= \int \left(\lambda''(t)\right)^2 dt.
\end{aligned}
$$

The quantity $R_{22}$ is analogous to the regularization quantity that `SMC++` uses, if specified (Terhorst et al., 2017).

We enforce these regularization constraints into our method by modifying the objective function during the M-step. The parameter update during EM, defined in equation (18) in the main text, then becomes

$$
\vec{\lambda}^{k+1} = \text{Argmax}_{\vec{\lambda}} \left[ Q(\vec{\lambda} | \{d_i\}, \vec{\lambda}^k) - \sum_i c_i R_i \right],
$$

where $i$ spans the various regularization types and $c_i$ are the respective weights used to adjust their relative strengths. The user can forgo the option to include any regularization by simply selecting 0 for the coefficients (the default behavior of our software implementation).

We note that the above definitions of the regularization quantities can be readily computed when the coalescent rate functions are continuous (as in the spline representation), however for the default piecewise constant representation of $\lambda(t)$ we use the generalized notion of derivatives and integrals for discrete domains instead. The derivatives are computed from finite differences between the constant values, and integration becomes summation over the constant pieces.

To demonstrate the impact of regularization on the inference using our method, we re-analyzed the data simulated under the population history with a bottleneck followed by exponential growth presented in Figure 9 in the main text, and vary the strength of regularization. In particular, we analyzed the simulated datasets with 10 haplotypes using the composite model CHIMP-$\mathcal{T}_{2,5,10}$ with the same piecewise constant parametrization using 20 pieces, as before. We set all regularization coefficients $c_i$ to zero, except for $c_{12}$, the coefficient controlling $R_{12}(\lambda(t))$ (squared first derivative), which we varied over several orders of magnitude, specifically, $c_{12} \in \{10^{-7}, 10^{-6}, 10^{-5}, 10^{-4}, 10^{-3}\}$, and depict the results in Figure A. We observe that for very strong regularization, $R_{12}(\lambda(t))$ is forced to be close to zero, and the inferred history is nearly constant, whereas for weak regularization, the performance is similar to the unregularized case depicted in Figure 9 in the main text. For intermediate regularization, the inferred histories show varying degrees of smoothness.

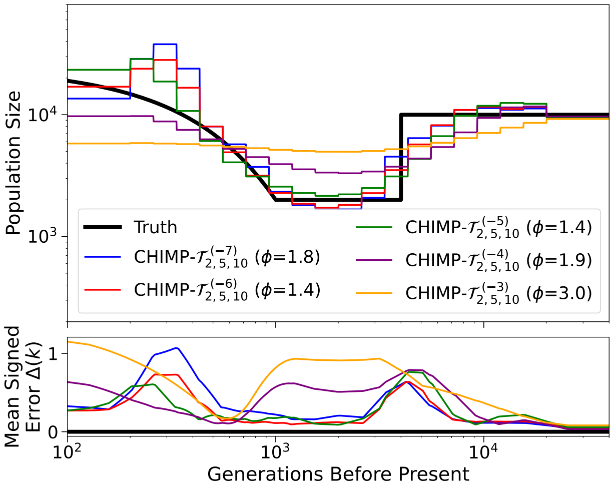


Figure A: CHIMP-$\mathcal{T}_{2,5,10}$ is used to infer the population size history from 10 simulated haplotypes in the scenario of a bottleneck followed by exponential growth. The regularization coefficient $c_{12}$ takes values $\{10^{-7}, 10^{-6}, 10^{-5}, 10^{-4}, 10^{-3}\}$, and the respective exponent of 10 is indicated in the superscript in the legend.

### 3 Choosing Interval Boundaries for Hidden States

For both $T_{MRCA}$ and $\mathcal{L}$, we have the task of deciding how to partition the continuous values for these quantities into discrete states to use for the CHMM. For $T_{MRCA}$, we follow Li and Durbin (2011) and Wang et al. (2021), and divide the continuum into roughly equidistant intervals on an exponential scale. Specifically, to divide the positive real line into $S$ intervals for the hidden states, we choose the interval boundaries

$$t_i := \alpha * \big(\exp(i/(S-1) \cdot \log(1 + t_{\max}/\alpha)) - 1\big), \tag{S1.1}$$

for $i \in \{0, \ldots S\}$ and $t_S = \infty$, with $\alpha = 0.01$ and $t_{\max} = 25$. Note that $t_0 = 0$, $t_{S-1} = t_{\max}$, and the increments from $t_i$ to $t_{i+1}$ get larger with increasing $i$. We experimented with adjusting these intervals for different subset sizes $n_s$ in the composite likelihood, but we found that the unadjusted partitioning worked well across the different $n_s$ tested here.

For the CHMM using the tree length $\mathcal{L}$ as hidden state, we choose a different strategy and divide the continuum into states such that the marginal probabilities of the CHMM occupying any one state at a given locus are approximately uniform. We compute this partition for the uniform distribution in a model of constant population size, where we estimate the size from the number of segregating sites in the given sample using Watterson's estimator (fixing the per generation mutation rate). In order to partition the continuum, we numerically partition it so that the distribution of $\mathcal{L}$ is uniform across the states. The analytical formulas for the distribution of $\mathcal{L}$ in the case of a constant population size is

$$f_{\mathcal{L}}(t) = \frac{n-1}{2} e^{-t/2} \left(1 - e^{-t/2}\right)^{n-2},$$

see for example equation (3.34) of Wakeley (2009). By numerically evaluating these distributions, we find the respective uniform partition.

We experimented with using a corresponding uniform distribution for $T_{\mathrm{MRCA}}$, but found that the partition detailed in equation (S1.1) resulted in more accurate inference. We do not have analytic results explaining why it performs better, but it appears necessary that the partition has several discretization points more ancient than the last changepoint of the population size history. This often results in some states on the fringes having very low mass in the marginal distribution. Similarly, we tried using an adjusted discretization like equation (S1.1) for the CHMM with $\mathcal{L}$ as the hidden state, but the numerical solutions of the PDE became more unstable, since some states would have very little mass.

Since the computing time required for the forward-backward algorithm scales quadratically with the number of hidden states (matrix multiplication), even using 50 states may create a computational bottleneck. However, using the computational improvements detailed in Section 4 can compensate and make the algorithm tractable again by reducing the number of times matrix multiplication is performed.

### 4 Improving Speed for Forward and backward Algorithm

Computing the entries of the full forward and backward table for each nucleotide site of the genome using the respective algorithm would be highly inefficient, and thus we implement two different ways to improve here. The *locus-skipping* efficiently integrates over large monomorphic regions without segregating sites, whereas the *meta-locus model* approximates the likelihood to reduce the number of "effective" loci for the CHMM.

#### 4.1 Locus-Skipping

The *locus-skipping* method to improve efficiency was implemented as described in the supplement of Terhorst et al. (2017). This is a modification to the forward and backward algorithm that computes the exact likelihood for tracts of monomorphic sites in a single-step, thereby greatly improving efficiency. Briefly, this method uses the eigendecomposition of the single-step transition matrix at monomorphic sites to efficiently exponentiate the respective matrix and integrate large tracts of monomorphic sites in a single step. It is important to note that the underlying likelihood model is exact and equal to the model that considers each site individually. Thus, it retains all the relevant information necessary to compute the expectations required in the EM algorithm.

We also note that for the locus-skipping step some of the probabilities are stored as *log* values to avoid machine precision issues. Even with this measure numerical errors arise when the tract being skipped is too large, and in practice we cap the size of these skips at 1000 loci. Tracts of missing data are straightforward to handle in this case as well, since can be treated as monomorphic tracts with modified emission probabilities, where each hidden state is equally likely to emit a missing nucleotide.

#### 4.2 Meta-locus Model

The *meta-locus* model is an alternative to the *locus-skipping*, and involves a slight modification of the underlying likelihood model itself, following a strategy that has been described in (Steinrücken et al., 2019, Suppl. Text 4.2). Rather than implementing the CHMM with a hidden state and an emission at each nucleotide site, we combine a specified number of sites into a meta-locus, with a single hidden state but with several emissions for all nucleotides that are grouped into this meta-locus. We specify in advance the

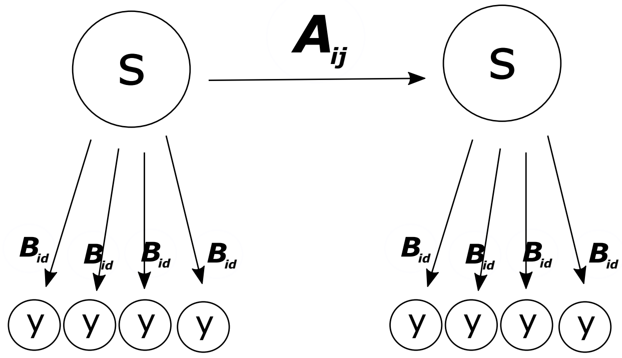


Figure B: Schematic for *meta-locus* model with $s = 4$. Each meta-locus represents 4 sites. Recombination does not happen within a meta-locus, but between at elevated rates. The emission probabilities $\mathbf{B}_{id}$, 4 for each meta-locus, are the same as in the single-step model.

number of nucleotide sites $s$ that a meta-locus spans, which also determines the number of emissions per locus. Figure B shows a schematic of this approximation.

The emission probabilities for each site in this model are identical to those used in the full model. However, by assuming a single hidden state for all sites in a meta-locus, we effectively suppress recombination between these, and thus it is necessary to implement recombination between two meta-loci at correspondingly elevated rates. To this end, the transition probabilities are obtained by raising the single-site transition matrix to the power $s$ (the number of sites in the meta-locus).

This implementation offers the potential for a tremendous speedup of the forward-backward algorithm and this option should be used used when the E-step appears to be the computational bottleneck during inference, in particular when the number of discrete CHMM states is large.

### 5 Data Processing

Our algorithm can be applied to full genomic sequencing data for a sample of individuals from a population. Such data are often presented as genetic variation at a number of segregating sites, separated by tracts of monomorphic sites, for example, in the form of a `vcf`-file. Additionally, such datasets often have regions in the genome marked as missing. This suggests that all the sites along the genome can be grouped into tracts of two types: *non-segregating* tracts, defined as having a single segregating bi-allelic site at the beginning, followed by a stretch of non-segregating sites, and *missing* tracts, where the genetic information is flagged as missing in the data. Processing the raw data and opting to store it as a series of tracts is more efficient than storing information for each individual site, and by also storing the type of tract, the tract length, and the number of derived alleles at the head of *non-segregating* tracts, we retain all the necessary information to provide to our CHMM. Lastly, our method needs information on whether a certain allele at a segregating site is ancestral or derived, which can be specified by providing an ancestral sequence in addition to the population sample.

#### 5.1 Missing Data

##### 5.1.1 Non-segregating sites in CHMM

While the *non-segregating* tracts can be handled straightforwardly by the CHMM, with the monomorphic non-segregating sites being treated as an emission of 0 derived alleles at the respective site, the missing tracts are handled slightly different. If a site is missing, the emission probabilities that are used for that site during the forward-backward algorithm are modified to reflect the fact that the missing site provides no information on the hidden state at that location. In practice, this means that each hidden state has an

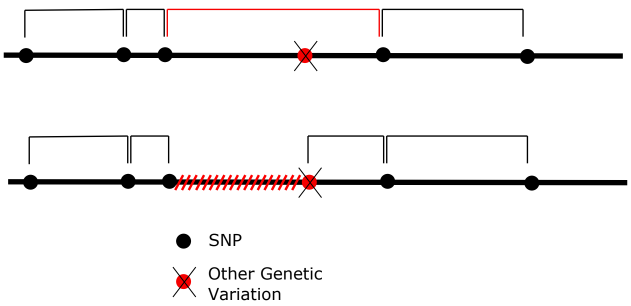


Figure C: This schematic shows the difference between just tagging a non bi-allelic segregating site as missing and also masking the tract upstream as missing. In the former case (top), the tract is effectively interpreted as an artificially long region without variation (marked red), which biases the inference. By masking the left region as missing (bottom) we avoid this systematic bias.

equal probability of emitting a missing site. For tracts of missing data, this has the effect that the posterior probability distribution across hidden states at the boundaries is informed by the surrounding sequence, but further into the interior of such tracts the posterior probability approaches the marginal distribution of states asymptotically.

##### 5.1.2 Tagging

We note that genomic datasets can contain segregating sites that are not bi-allelic, for example, segregating sites with 3 alleles, or some forms of structural variation. We treat all segregating sites that are not bi-allelic as *missing* sites. This step introduces a potential bias into the CHMM because it can potentially flag individuals segregating sites as missing, while the non-segregating sites around this specific sites are still used for the analysis. A single missing site is more likely to be non-segregating a-priori, and if such a site is between two tracts of non-segregating sites, the method effectively concatenates them and interprets the whole tract as an artificially long tract with no segregating sites. This biases the CHMM towards inferring shallow trees. In order to counter this systematic bias, when we tag such segregating sites that are not bi-allelic as *missing*, we also mask all the non-segregating sites upstream as missing, until we reach either a segregating site or a site marked as missing in the raw data (see Figure C for a schematic). By marking the tract upstream (or equivalently downstream) as missing as well, the true non-segregating tracts that are identified better represent the appropriate lengths of such tracts. At worst, this process loses information since the inferred trees in missing tracts approach the marginal distribution asymptotically as the algorithm moves into the interior of these tracts, but it does not introduce systematic bias.

#### 6 Heuristics for Discretization of Population Size History

Figure D shows the results when inferring the population size history in the bottleneck followed by exponential growth scenario (see Section 3.3 in the main text), when using the default discretization for the different methods. For the method `CHIMP`, the default implementation chooses the discretization as follows: first we compute $\hat{N}$ from Watterson's estimator, and then partition the interval $[\frac{\hat{N}}{50}, 20 \times \hat{N}]$ (in generations) into 18 logarithmically equidistant epochs with an additional epoch being added above and below the minimum and maximum. Choosing an inappropriate discretization can have an impact on the accuracy of the inference, and can also lead to the inference missing important features of the underlying history.

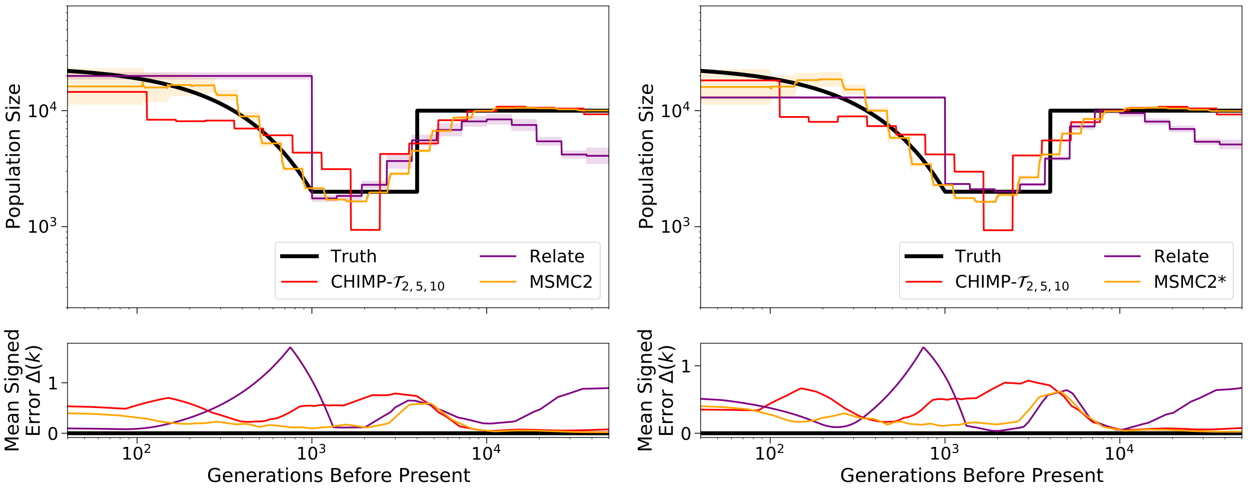


(i) Sample size 10.

(ii) Sample size 200.

Figure D: Results of inference in the bottleneck followed by growth scenario for sample size (i) 10 and (ii) 200. We compare the results of CHIMP-$T_{\mathrm{MRCA}}$, MSMC2, and Relate using the respective default discretization of the population size history. Truth shown in black. Solid lines are averages over 16 replicates and shaded area indicates standard deviation. Mean signed error $\Delta(k)$ is shown in bottom plot and has been smoothed using moving average for visualization purposes and the integral $\phi$ is indicated in the legend. (*) For sample size 200, MSMC2 was run on 50 non-overlapping pairs.

### 7 Demographic Models with Limited Number of Parameters

We also compared the inference methods on two simple demographic models with three parameters to estimate. The first models a population that experienced a bottleneck event. In this scenario, the ancestral population is of constant diploid size 10,000. At 2,000 generations before present, the population size is reduced to 5,000, but recovers to 10,000 at 1,000 generations before present. The second scenario models a population experiencing piecewise growth. Here, the ancestral population size is again 10,000 diploid individuals. At 2,000 generations before present, the population size doubles to 20,000, and then doubles again at 1,000 before present to 40,000. In both scenarios, we fixed the demographic parameterization for each method to estimate a three-parameter piecewise constant population size history with change points matching those of the true population size histories.

Figure E shows the results of the inference using the different methods in the bottleneck scenario for sample sizes 10 and 200. In this case, all versions of CHIMP and MSMC2 recover the true population size history accurately and show little variability across replicates. Relate, however, does not recover the underlying true size accurately. For a sample of size 10, it does not infer a bottleneck, and while for sample size 200, a bottleneck is inferred, the ancestral and contemporary population size are not inferred correctly. Nonetheless, the inference shows little variability.

The results for the inference in the piecewise growth scenario are depicted in Figure F. In this scenario, all versions of CHIMP are again able to infer the population size history with high accuracy. For a sample of size 10, MSMC2 also recovers the true size, however, for samples size 200, the inference for the intermediate size is systematically biased, with little variability. The reason for this deviation is likely the fact that when using MSMSC2 for large sample sizes, the interface does not allow setting the change points for the inference closer to the true values. Given this constraint, the inferred population sizes are quite accurate and we believe that the method would have high accuracy if the appropriate boundaries could be specified. Again, for a small sample, Relate overestimates especially the intermediate size, and for the large sample size of

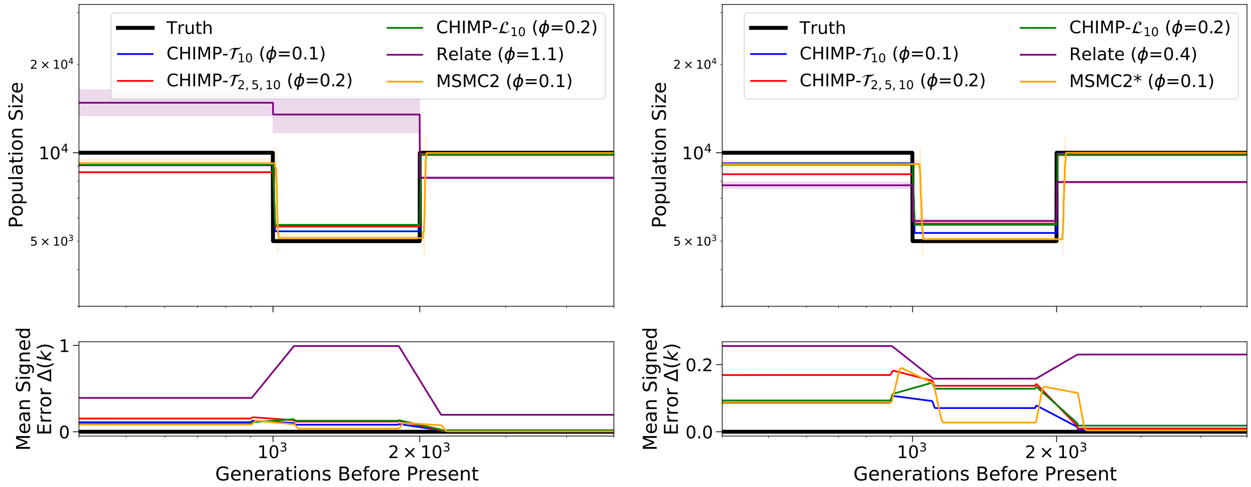


(i) Sample size 10. (ii) Sample size 200.

Figure E: Results of inference in the bottleneck scenario for sample size (i) 10 and (ii) 200. We compare the results of `CHIMP`, `MSMC2`, and `Relate` to infer the three population sizes, fixing the change points to match the truth (shown in black). Solid lines are averages over 16 replicates and shaded area indicates standard deviation. Mean signed error $\Delta(k)$ is shown at bottom and has been smoothed using moving average for visualization purposes. The integral $\phi$ is indicated in the legend. (*) Note that for sample size 200, `MSMC2` was only run on 50 non-overlapping pairs.

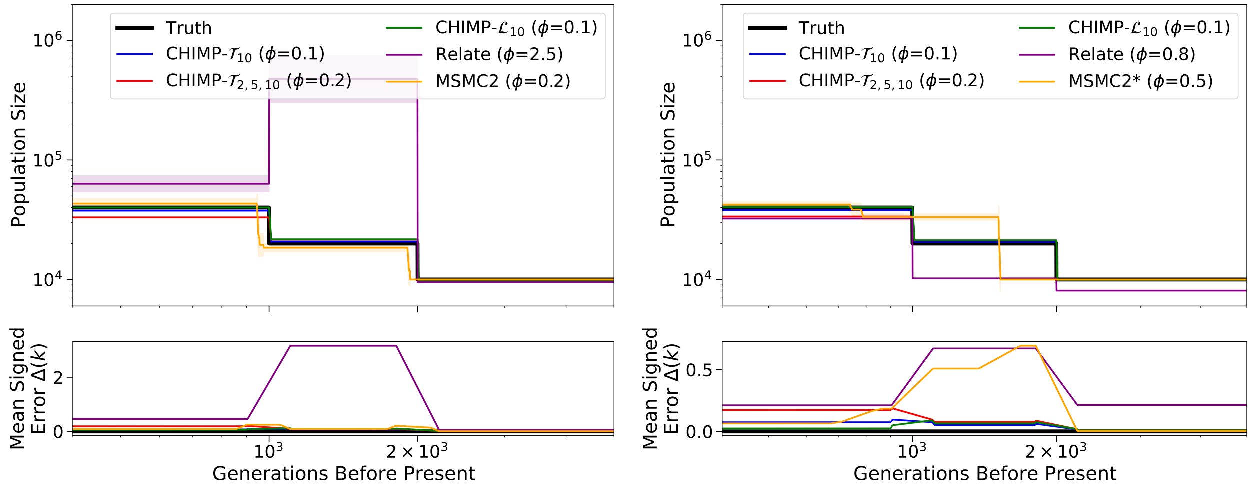


(i) Sample size 10. (ii) Sample size 200.

Figure F: Results of inference in the piecewise growth scenario for sample size (i) 10 and (ii) 200. We compare the results of `CHIMP`, `MSMC2`, and `Relate` to infer the 3 population sizes, fixing the change points to match the truth (shown in black). Solid lines are averages over 16 replicates and shaded area indicates standard deviation. Mean signed error $\Delta(k)$ is shown in bottom plot and has been smoothed using moving average for visualization purposes. The integral $\phi$ is indicated in the legend. (*) Note that for sample size 200, `MSMC2` was only run on 50 non-overlapping pairs.

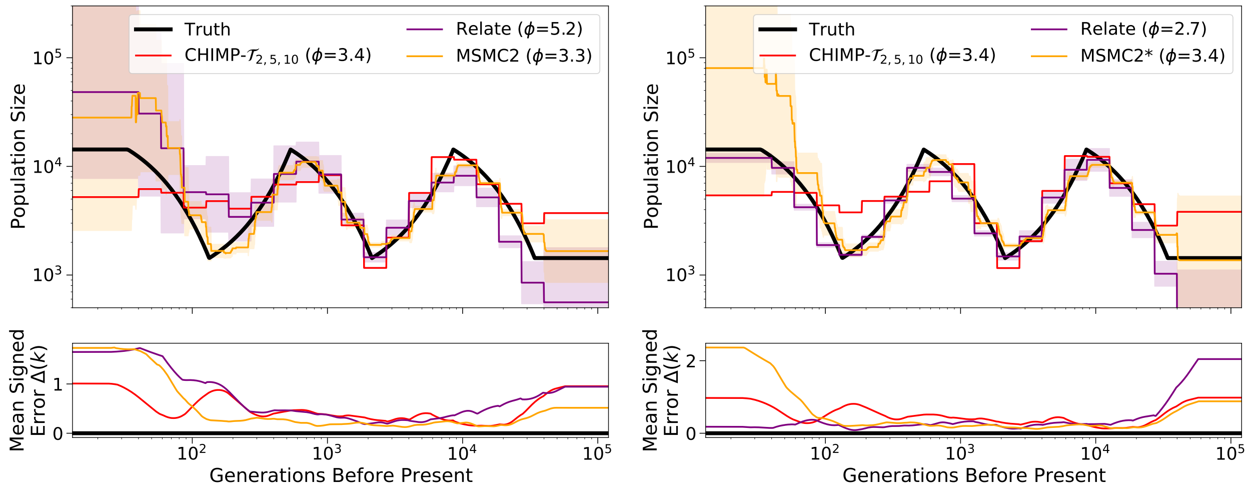


(i) Sample size 10.

(ii) Sample size 200.

Figure G: Results of inference for the sawtooth demography simulated using a human recombination map (for Chromosome 3) for sample size (i) 10 and (ii) 200. We compare the results of CHIMP-$T_{\mathrm{MRCA}}$, MSMC2, and Relate. For Relate we provide the recombination map used to simulate the data, for CHIMP we specify the constant recombination as the mean over the recombination map, and MSMC2 is set to infer the best fitting constant rate. Truth shown in black. Solid lines are averages over 16 replicates and shaded areas indicate standard deviation. Mean signed error $\Delta(k)$ is shown at bottom plot and has been smoothed using moving average for visualization purposes. The integral $\phi$ is indicated in the legend. (*) For sample size 200, MSMC2 was run on 50 non-overlapping pairs. Only 15 out of 16 replicates finished for MSMC2.

200 the estimates are not as accurate as MSMC2 and CHIMP.

### 8 Simulations with Recombination Map

Here, we explore the effects of varying recombination rates along the chromosome on demographic inference using the different methods. We used msprime to simulate data under the *sawtooth* demography (explored in Section 3.3 in the main text) using the HapMap II recombination rates on Chromosome 3 (The International HapMap Consortium, 2007). We inferred the population size history using CHIMP-$T_{\mathrm{MRCA}}$, MSMC2, and Relate, and compared their performance using a the same discretization that was used in Figure 8 in the main text. We provided the methods with as much information about the recombination rates as possible: we provided Relate with the recombination map used to simulate the data. For CHIMP-$T_{\mathrm{MRCA}}$, we specified a constant recombination rate corresponding to the mean recombination rate. Lastly, MSMC2 was set to infer a recombination rate while optimizing the population size history.

The results of these inferences are shown in Figure G. Comparing these results to those in Figure 8 in the main text, we note that CHIMP-$T_{\mathrm{MRCA}}$ and MSMC2, while operating under the assumption of a constant recombination rate, perform similarly well as in Figure 8 in the main text, where the recombination rate was in fact constant. We also note that Relate, especially for 200 haplotypes, performs much better in the case of varying recombination rates. The reason for this improvement is likely the fact that the procedure used in Relate to infer genealogical trees can be shown to infer the correct tree in regions without recombination. Since there are many *cold* spots in the human recombination map, Relate infers the true underlying genealogies more accurately in these regions, and this yields an overall improvement in performance.
